# Trait-environment relationship in tadpoles of the southern Atlantic Forest

**DOI:** 10.1101/2021.10.08.463663

**Authors:** Roseli Coelho dos Santos, Diego Brum, Diego Dalmolin, Renata Krentz Farina, Elaine Maria Lucas, Alexandro Marques Tozetti

## Abstract

1. Environmental predictors select individuals by their functional traits, shaping the anuran assembly patterns. Individuals respond to environmental filters that can be on a local or regional scale.
2. In this study, we investigated the association between local (water and microhabitat) and landscape variables and the morphological traits of tadpoles of ponds and streams. The study was conducted in the southern region of the Brazilian Atlantic Forest. We sampled 28 waterbodies and recorded 22 anurans species. We performed RLQ and fourth-corner analyses to determine the patterns of trait-environment relationships and determine which environmental and landscape variables influence the morphological characteristics of tadpoles from streams and ponds.
3. We found that the morphological traits of tadpoles are influenced mainly by physicochemical and microhabitat attributes, being distinct between ponds and streams. In ponds, water depth, pH, and the presence of vegetation influence the morphological traits of the tadpoles, while in the streams water pH, temperature, conductivity, total alkalinity, Alk HCO3, and microhabitat variables played a major role in defining the traits.
4. Our results indicate that local components of habitat (water characteristics and microhabitat) influence functional traits of tadpoles in both ponds and streams, especially those supposedly related to locomotory, foraging and prey-detection abilities.

## INTRODUCTION

Functional diversity is a biodiversity measurement based on functional traits of the species present in a community and is evaluated by measuring the value and variation of functional characteristics that are prevalent in the ecosystem (Goswami, Bhattacharyya, Mukher, & Tribedi, 2017). It aims to analyze how and how much an organism’s traits influence its performance, reflecting on the functioning of the ecosystem (Diaz & Cabido, 2001). The functional diversity can be associated with morphological, physiological and phenological variations at the individual level, as well as with ecosystem processes and patterns (Petchey, O’Gorman, & Flynn, 2009). Functional traits are the main determinants of the biology of organisms through biochemical, physiological, morphological, developmental or behavioral mechanisms (Violle, Navas, Vile, Kazakou, Fortunel, Hummel, & Garnier, 2007). Functional diversity studies have been applied to analyze the trait-environment relationship in several taxa, including plants (Liu, Swenson, Lin, Mi, Umaña, Schmid, & Ma, 2016), fishes (Carvalho & Tejerina-Garro, 2015), birds (Arruda Almeida, Green, Sebastián-González, & Dos Anjos, 2018) and amphibians (Lescano, Miloch, & Leynaud, 2018; Jordani, Mouquet, Casatti, Menin, Rossa-Feres, & Albert, 2019; Dalmolin, Tozetti, & Pereira, 2020; Lipinski, Schuch, & Santos, 2020).

Due to their biphasic life cycle and their water-dependent physiological characteristics, amphibians form a suitable group for studies of functional diversity. Tadpoles occur in a variety of freshwater habitats, including ponds, streams, phytotelmata of bromeliads and logs, as well as permanent or temporary shallow water-covered surfaces (Altig & McDiarmid, 1999). This adaptation to different habitats can result in great variation in ecological and morphofunctional characteristics of tadpoles, which are influenced by different selective pressures of the environment (Duellman & Trueb, 1994; Sherratt, Vidal-García, Anstis, & Keogh, 2017; Sherratt, Anstis, & Keogh, 2018). The different selective pressures act as environmental filters, selecting the species by their morphological traits (Violle, Enquist, McGill, Jiang, Albert, Hulshof, Vincent, & Messier, 2012). The environmental filters can act at different scales, such as micro-spatial (local habitat characteristics) and macro-spatial (landscape characteristics), determining the patterns of community assembly (Weiher, Freund, Bunton, Stefanski, Lee, & Bentivenga, 2011; Violle et al., 2012).

Studies with tadpoles indicate that some species show phenotypic or behavior plasticity to adjust their occurrence according to the characteristics of the environment (Nomura, Do Prado, Da Silva, Borges, Dias, & Rossa-Feres, 2011; Groner, Buck, Gervasi, Blaustein, Reinert, Rollins-Smith, Bier, Hempel, & Relyea, 2013; Laufer, Vaira, Pereyra, & Akmentins, 2015). Recent studies with tadpoles have turned their attention to functional diversity (Jordani et al., 2019; Dalmolin et al., 2020; Lipinski et al., 2020) and shown that environmental and spatial descriptors are determinants for taxonomic and functional composition. The functional morphology of tadpoles can be influenced by both local heterogeneity or soil use at the microhabitat level and landscape at the regional level (Queiroz, Silva, & Rossa-Feres, 2015; Marques, Rattis, & Nomura, 2018). Most studies of the functional diversity of tadpoles are based on taxonomic and phylogenetic (Violle et al., 2012; Jordani et al., 2019; Lipinski et al., 2020) and metamorphosis traits (Mogali, Shanbhag, & Saidapur, 2021), and few studies analyze tadpole morphological traits as a response to environmental factors, especially in tropical regions (Queiroz et al., 2015). However, identifying how morphological traits respond to local environmental and landscape characteristics can provide insight into the morphological traits of tadpoles that are crucial for community assembly, mainly in the southern portion of the Atlantic Forest, which presents a considerable anuran diversity.

On other hand, as tadpoles occur in a variety of environments with different hydrological regimes (e.g., streams and ponds), they are exposed to several selective pressures, showing ecological and morphofunctional variations ensuring their survival (Roelants, Haas, & Bossuyt, 2011; Jordani, Melo, Queiroz, Rossa-Feres, & Garey, 2017; Sherratt et al., 2017; 2018). Landscape changes are a focus of study by the scientific community due to the impacts caused on the structure and dynamics of ecosystems (Haddad, Brudvig, Clobert, Davies, Gonzalez, Holt, & Lovejoy, 2015). Environmental changes (e.g., land use, nutrient availability and cycling, atmospheric composition, climate, introduction of exotic species, and overexploitation by humans) have been promoting modifications of species diversity and composition (Hooper, Chapin, Ewel, Hector, Inchausti, et al., 2005). Species richness, composition, and abundance distribution, which are the basic dataset for ecology, are used as parameters to describe ecological community structure (Gotelli & Colwell, 2001), while functional diversity, based on the value and range of biological traits in ecosystems (Diaz & Cabido, 2001), has a greater impact on ecosystems process than on species diversity (Tilman, Knops, Wedin, Reich, Ritchie, & Siemann, 1997; Petchey, Hector, & Gaston, 2004; Hooper et al., 2005). Therefore, studies analyzing the functional diversity traits of tadpoles may help predict the impacts of these landscape and environmental changes on the structure of biological communities

In this sense, environmental factors, including morphological characteristics of microhabitat and physicochemical characteristics of the water, could be important to explain the functional diversity of tadpoles. Such environmental characteristics can directly influence the composition of aquatic communities in response to predation and/or competition (Wellborn, Skelly, & Werner, 1996; Werner, Yurewicz, Skelly, & Relyea, 2007; Melo, Garey, & Rossa-Feres, 2018), as well as nutrient supply (Williams, Rittenhouse, & Semlitsch, 2008), thus enabling the distribution of trophic guilds in the water column (Queiroz et al., 2015). They can also influence the metabolism (Afonso & Eterovick, 2007), morphology and physiology of tadpoles (Sipaúba-Tavares, Guariglia, & Braga, 2007; Thomaz & Cunha, 2010; Mansano, Stéfani, Pereira, & Macente, 2012; Mansano, Stéfani, Pereira, Nascimento, & Macente, 2014; Farquharson, Wepener, & Smit, 2016), shaping their performance. This prediction is based on the influence of the landscape on the composition of the tadpole community (Santos et al., in press) and the local characteristics of the water such as depth, presence of vegetation (Queiroz et al., 2015), substrate (Schiesari, 2006; Williams et al., 2008; Thomaz & Cunha, 2010), temperature (Lima, Casali, & Agostinho, 2003; Maciel & Juncá, 2009), pH, nitrogen in the form of ammonia, nitrite and nitrate, phosphorus and bicarbonate (Mansano, Vanzela, Américo-Pinheiro, Macente, Khan, et al., 2018).

This study aimed to investigate patterns of the trait-environment relationship in tadpole communities from streams and ponds. Our first hypothesis is that the functional diversity traits of tadpoles are influenced by local descriptors such as the physicochemical characteristics of the water and microhabitat as well as by landscape characteristics. Besides that, we searched for differences in these relationships between ponds and streams. Our second hypothesis is that environmental filters act differently in streams and ponds, selecting species with similar morphological traits for each habitat type. Based on the fact that streams and ponds are distinct environments in terms of hydrological characteristics and structure of aquatic communities (Schriever & Lytle, 2016; Jordani et al., 2017) it is very likely that there are differences in the influence of environmental attributes on the functional diversity traits of tadpoles.

## METHODS

### Study site

We carried out the study in remnants of the Atlantic Forest in southern Brazil, between coordinates 22°30’ to 33°45’S (latitude) and 48°02’ to 57°40’W (longitude) (Figure 1). The landscape of the region is formed by Atlantic Forest and high-altitude grasslands (Veloso & Góes-Filho, 1982; SOS Mata Atlântica / INPE, 2008) interconnected by urban and rural areas with the presence of pastures and agricultural plantations (Ribeiro, Metzger, Martensen, Ponzoni, & Hirota, 2009; Pillar & Vélez, 2010). The climate is humid subtropical (IAP, 2004), with rainfall ranging from 1600 to 2200 mm per year (Alvares, Stape, Sentelhas, Gonçalves, & Sparovek, 2013). The wavy relief is formed by plateaus, plains and escarpments (IAP, 2004), with areas of steep slopes and embedded valleys (Santa Catarina, 1986). The altitude varies from 300 to 1200 m above sea level.

**FIGURE 1.**
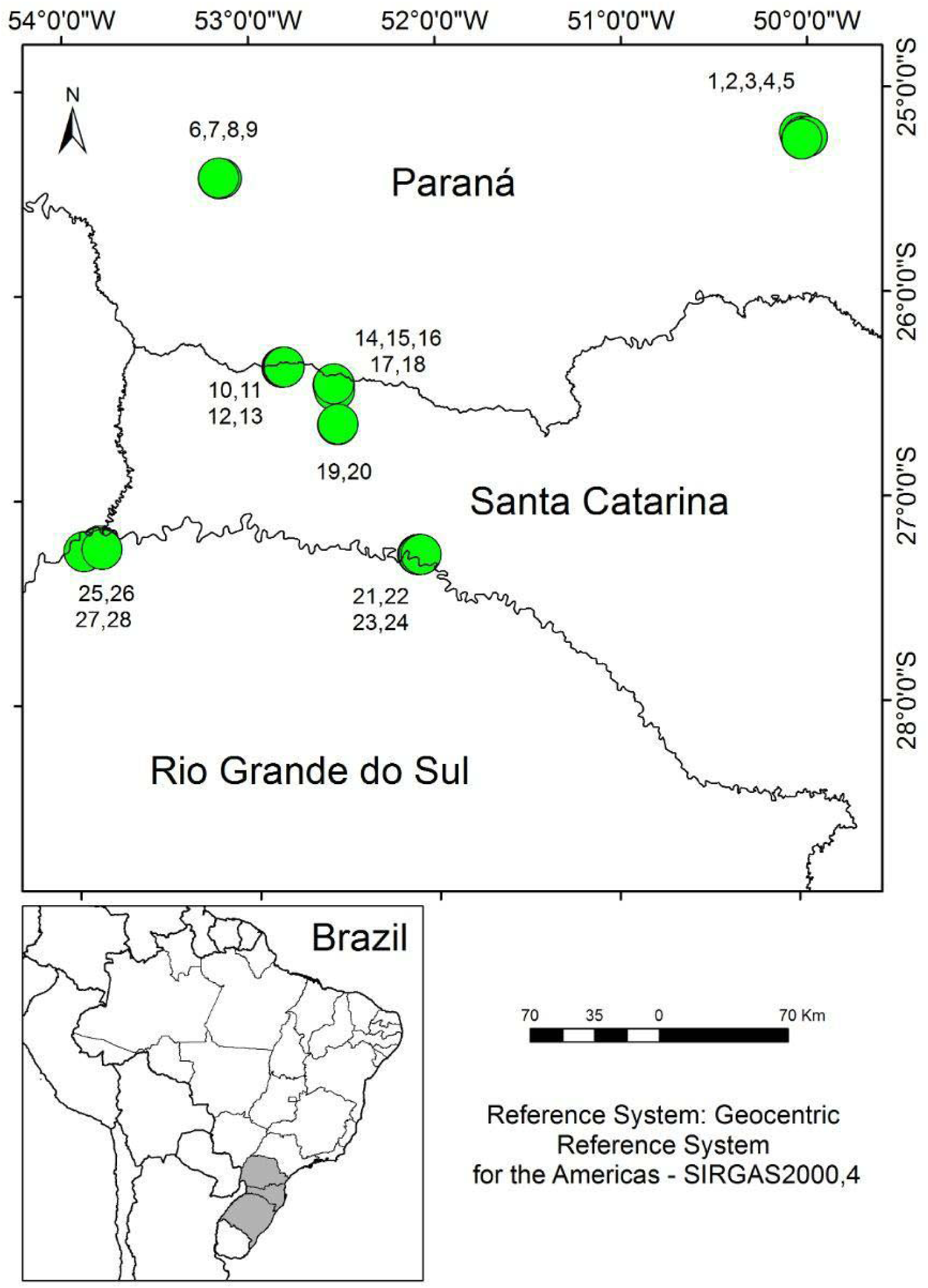
Sampled waterbodies in areas of Atlantic Forest in southern Brazil. Numbers (from 1 to 28) refer to each waterbody sampled. For additional details, see Supplementary information (Figure S1).

The selection of the sampling sites was based on the climatic, altitudinal and rainfall patterns and with a similar history of human interference. For this, we applied the following criteria for choosing sample areas: a) presence of well-preserved remnants of the Atlantic Forest (preferentially inside protected areas); b) similar climatic conditions of temperature and rainfall; c) minimum variation in altitude, which ranged from 300 to 900 m above the sea level; d) limited latitudinal amplitude, between 22°30 ‘and 33°45’S. Finally, the exact sampling point (waterbody), was defined based on prior knowledge of the occurrence of amphibians and of abiotic and biotic characteristics that indicate reproductive sites of anurans (shallow water, presence of floating vegetation, and abundant vegetation at the margins) (Maltchik, Rolon, Stenert, Machado, & Rocha, 2011; Knauth, Moreira, & Maltchik, 2018).

### Study design

We sampled tadpoles in 28 waterbodies (13 ponds and 15 streams), from October 2018 to March 2019. The waterbodies consisted of permanent and semi-permanent ponds and transects of streams up to 100 m that were associated with forest habitats. Samplings were standardized in two hours in each waterbody, within a single breeding season. This procedure was adopted considering a large sample area and the reproductive period of amphibians in warm seasons in southern Brazil.

We collected the tadpoles using a wired dip net with a mesh size of 3 mm and a diameter of 300 mm (Heyer, 1976). Collection in each waterbody took an hour and included the greatest variety of microenvironments along the margin of the ponds and streams (Vasconcelos & Rossa-Feres, 2005; Santos, Rossa-Feres, & Casatti, 2007; Both, Solé, Santos, & Cechin, 2009; Bolzan, Saccol, & Santos, 2016). We searched through the whole margin of the ponds and, for the streams, all possible locations were searched, covering approximately 100 m of linear margin (adapted from Jordani et al., 2017). Immediately after collection, the tadpoles were euthanized, following the Brazilian regulations for the use of animals (CONCEA, 2018), by immersion in a 2% lidocaine solution. Afterward, they were all individualized and stored in containers containing absolute ethanol. In the laboratory, we identified each specimen with the aid of a stereomicroscope and identification keys (e.g., Machado & Maltchik, 2007; Gonçalves, 2014). The collection and handling of specimens were authorized by ICMBio (# 64962), state agencies (IAP / PR # 33/2018, IMA / SC # 01/2019, SEMA / RS # 37/2018) and the Animal Ethics Committee of the University of Vale dos Rio dos Sinos (CEUA-UNISINOS # PPECEUA08.2018).

### Environment predictors of waterbodies

We considered 33 environmental variables for each waterbody (Table 1). Variables were classified as (1) local environmental descriptors (physicochemical characteristics of the water and microhabitat configuration) and (2) landscape environmental descriptors. Regarding local descriptors, we measured 12 physicochemical characteristics of the water and 16 microhabitat components. Regarding the landscape, we evaluated five variables. For a better understanding, all evaluated variables are listed in Table 1, followed by a brief explanation about their relevance in an ecological context.

**TABLE 1.**
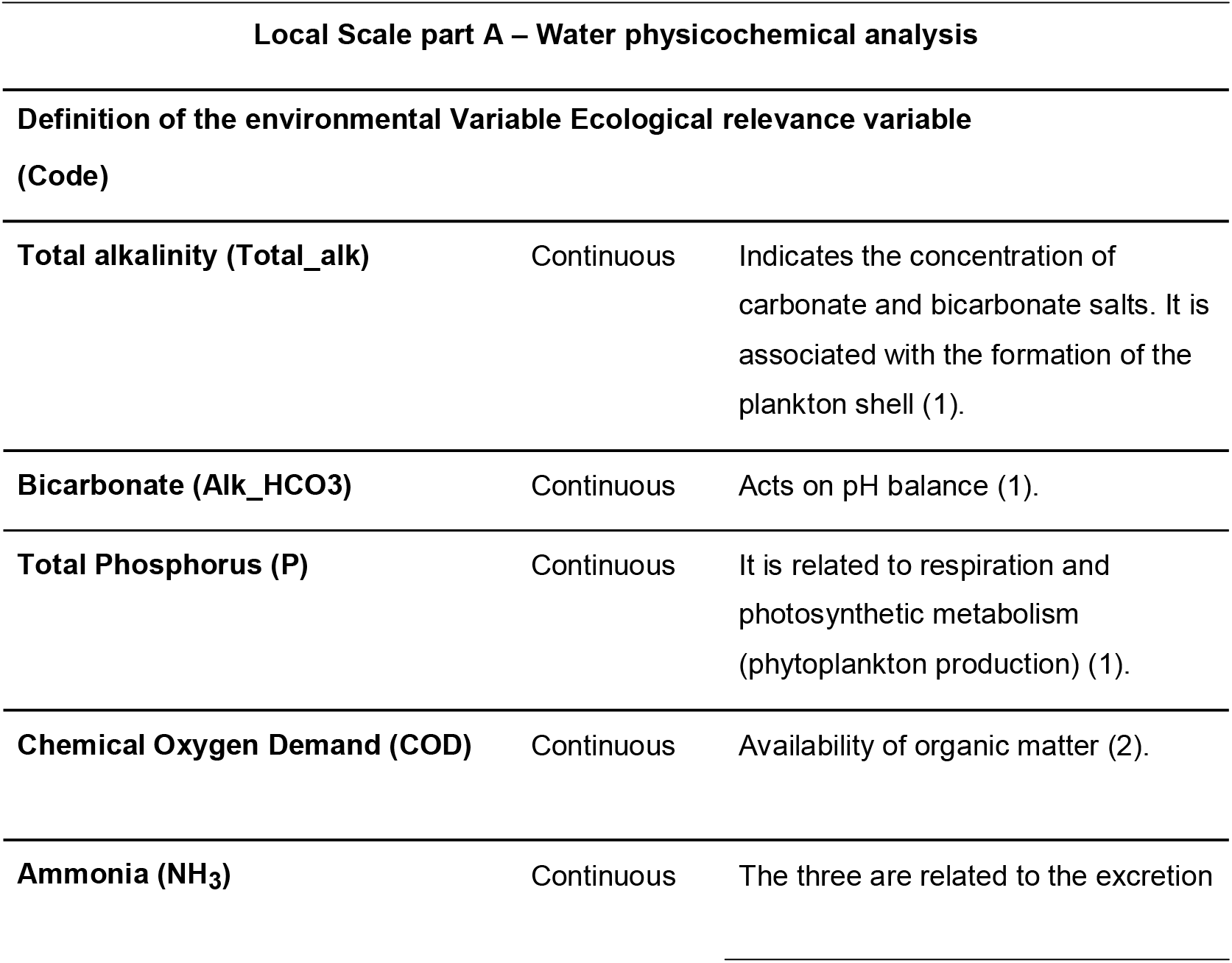
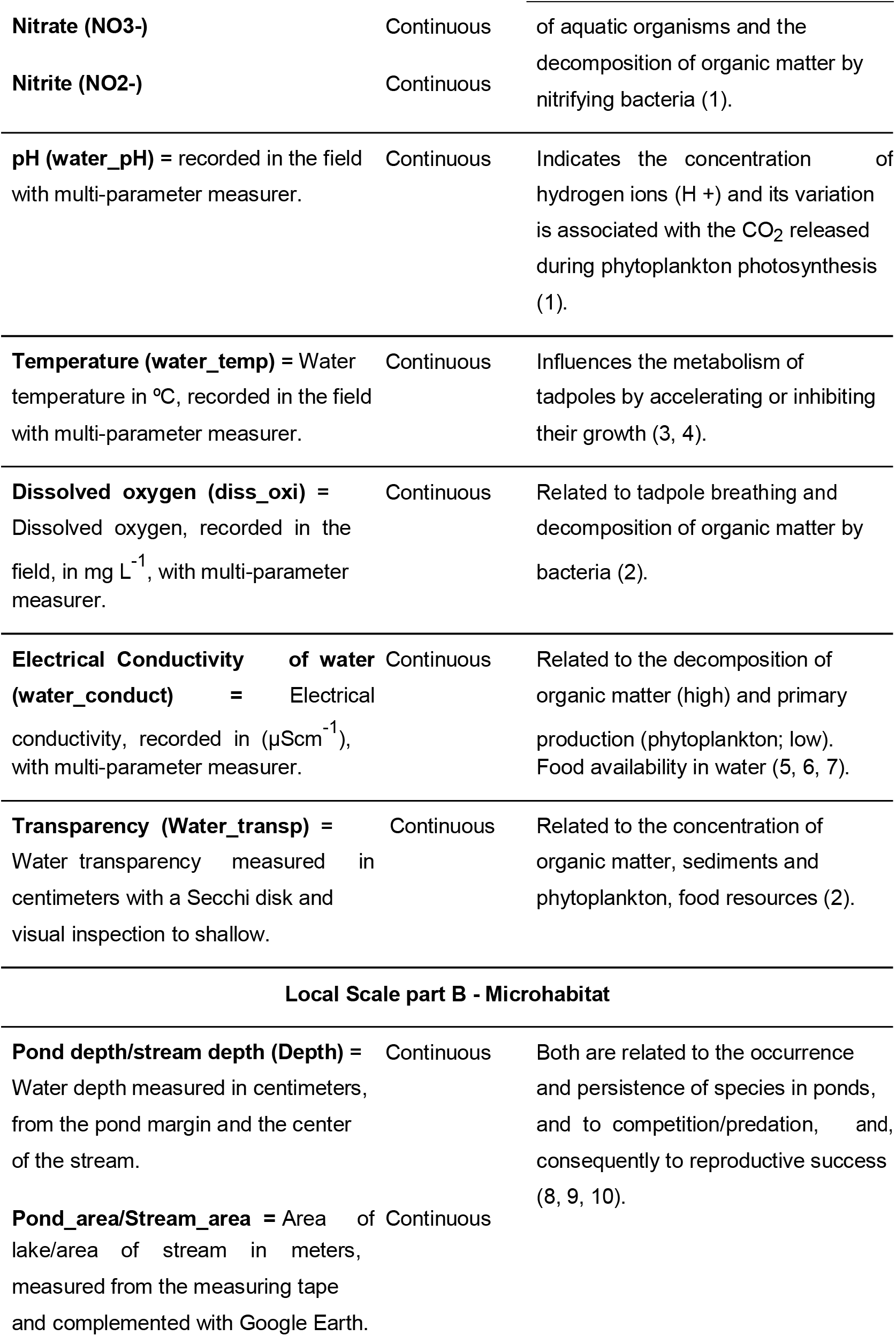
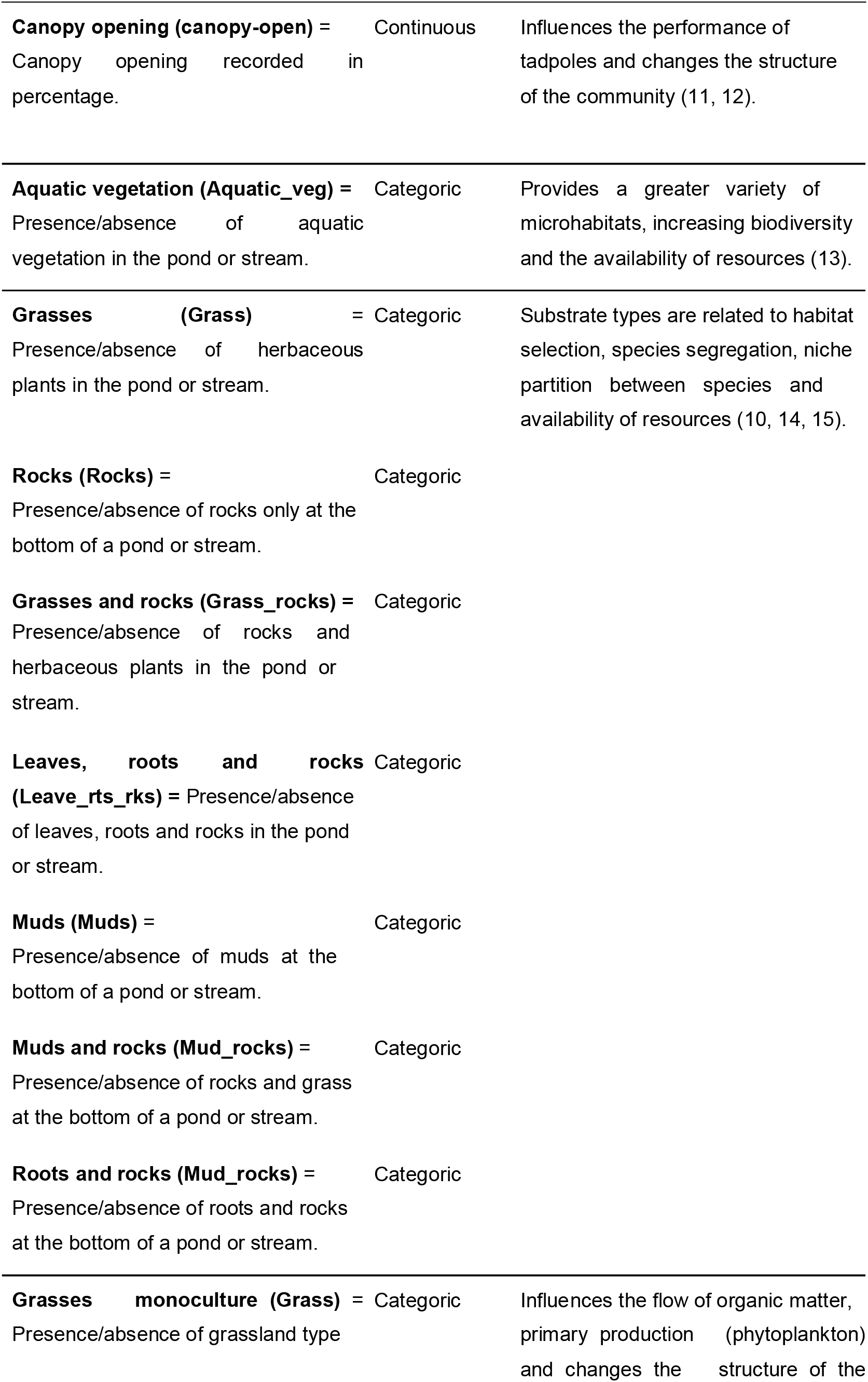
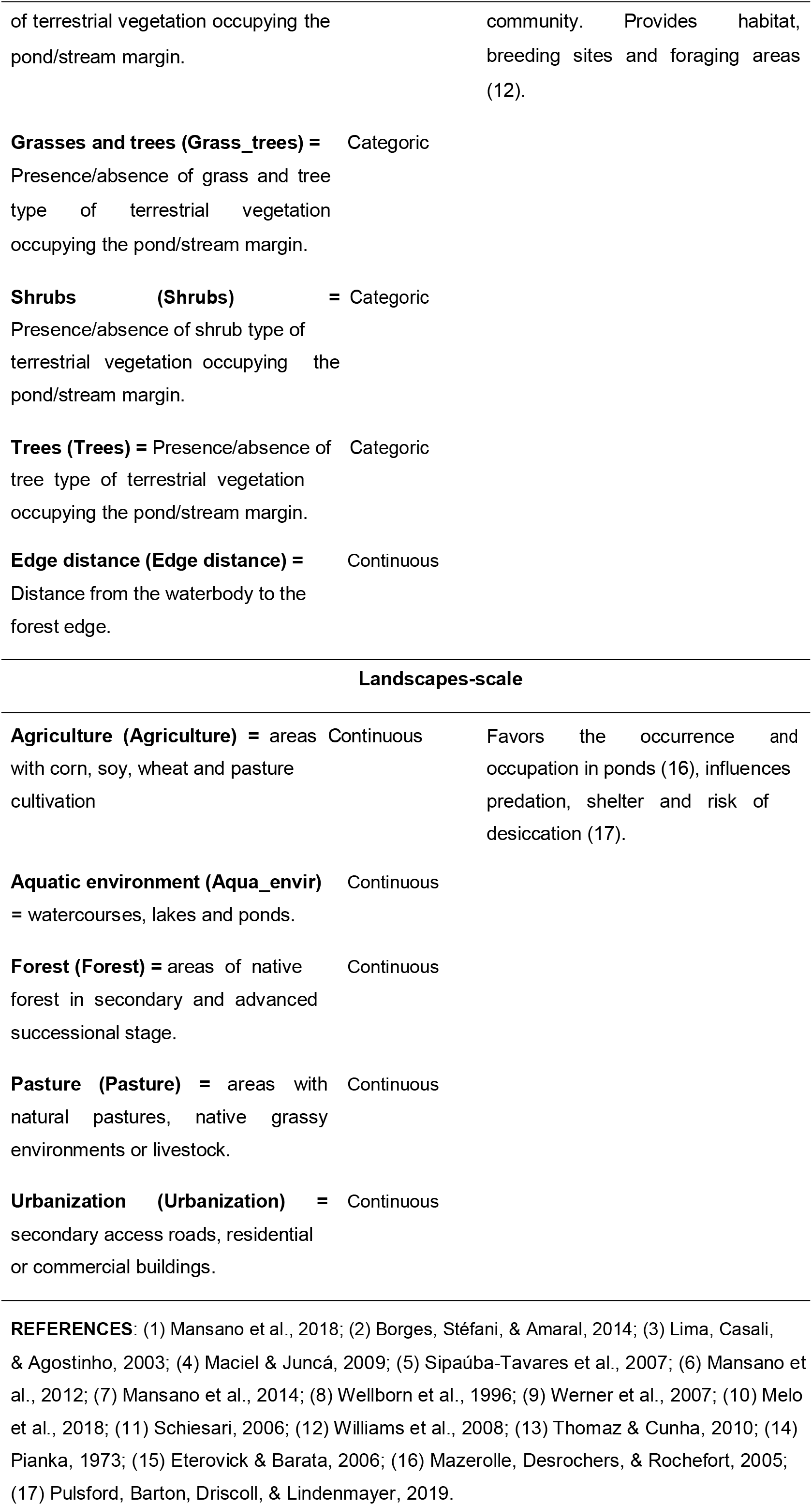
Environmental variables measured from sampled ponds and streams in tadpole collections. (A) Physicochemical – local scale, (B) Microhabitat – local scale and (C) Landscape-scale.

#### Local descriptors evaluation

For physicochemical analysis, we collected samples of surface water (15 cm deep), using sterilized dark plastic bottles (500 ml) that were later placed on ice. Samples were randomly collected at each tadpole collection site, 10 cm from the edge. The time between collection, transport and analysis did not exceed three days. In the laboratory, we conducted analyses of total alkalinity (Total_alk), bicarbonate (Alk_HCO3), total phosphorus (P), chemical oxygen demand (COD) and nitrogen in ammonia (NH3), nitrite (NO2-) and nitrate (NO3-) (APHA, 1998; Ternus, Souza-Franco, Anselmini, Mocellin, & Dal Magro, 2011). In the field, we collected data on pH (water_pH), temperature (water_temp) (°C), dissolved oxygen (diss_oxi) in mgL^-1^ and electrical conductivity (water_conduct) (µScm^-1^) using a multiparameter meter (*Lovibond Sensodirect* 150). Water transparency was measured with the Secchi disk, inserting approximately 15 cm deep. For shallower water bodies where it was not possible to apply the standard Secchi disk protocol, we visually estimated the transparency, recording its maximum viewing depth. The transparency was measured at the same tadpole-collection points and was done on the banks of the ponds and in the midpoint of the streams. All data were taken by the same observer to avoid skewed weightings. These variables are considered important for tadpoles since water quality has a direct influence on their behavior and development and an indirect influence on food availability (Castaneda, 2014; Zongo & Boussim, 2015).

For microhabitat data, we analyzed water depth, area, canopy opening, vegetation presence, and substrate inside and outside waterbodies. Pond and stream area was recorded using a measuring tape (in meters), and pond area was supplemented with polygon images (*Google Earth images*). For the ponds, we considered the area of the polygon and, for the area of the streams, we considered the width x the distance covered in each stream (maximum distance of 100 m). The area was presented in square meters (m^2^). Water depth was measured at the same points of tadpole collection and was recorded in centimeters, with records at three points for each waterbody. The images of the canopy opening were recorded using a spherical lens (*Universal clip lens*; 360°) coupled to a cell phone (*Xiaomi MI*) at the tadpole-collection point and later treated in the *GapLight* program version 2.0 (Frazer, Canham, & Lertzman, 1999) and presented as a percentage. For the canopy opening, we conducted a single measurement per water body, at the edge of the pond and in the center of the stream.

We recorded data of presence/absence on vegetation and substrate qualitatively by visual inspection. Initially, we recorded the presence of substrates inside water bodies and classified them into eight types: Aquatic vegetation (Aguatic_veg), Grasses (Grass), Rocks (Rocks), Grasses and rocks (Grass_rocks), Leaves, roots and rocks (Leave_rts_rks), Muds (Muds), Muds and rocks (Mud_rocks) and Roots and rocks (Roots_rocks). Following, we recorded and classified the waterbody’s terrestrial or marginal vegetation into four types: Grasses (Grass), Grasses and Trees (Grass_trees), Shrubs (Shrubs), and Trees (Trees). We considered phorophytes > 2 m as trees and < 2 m as shrubs, while grasses included native herbaceous plants (e.g., Cyperaceae, Poaceae and Typhaceae) and pasture monocultures (e.g., *Brachiaria* sp). For the qualitative recording of the substrate and terrestrial vegetation, we used four quadrants of 4 m^2^ in the interior and margins of the water bodies, with the margins of the ponds and the 100 m transects in the streams as the recording point. Finally, the distance from the forest edge was measured in meters using tape, taking as a starting point the edge of the pond/stream to the transition point of forest vegetation with roads and/or open areas.

### Landscape *descriptors evaluation*

We evaluated land use in the sampling sites based on the analysis of satellite images. We used Landsat 8 multispectral images, OLI sensor, available in the United States Geological Survey (USGS) and captured between January and July 2019. We selected images with minimal cloud coverage and without significant radiometric noise. We performed the following pre-processing steps of the images: 1) geometric corrections, due to the inherent geometric distortions that images collected at different times have, through the georeferencing of these images; 2) atmospheric corrections to reduce the interference of atmospheric scattering on the images (Soares, Almeida, Rubim, Barros, Cruz, et al., 2015); and 3) mosaic and enhancement of the different images in each season to reduce the seasonal effects on the visual aspect of the images. Pre-processing was done using the *ENVI* software, version 5.51. After the pre-processing steps, we defined classes of land use and occupation based on on-site observations, considering the predominant use of the areas. The categories were established as Agriculture, Aquatic environments, Forest, Pastures and Urbanization (Table 1). We classified the images based on their vectorization in ArcGIS software version 10.3, considering the buffer with a 250-m radius for each waterbody. The buffer size was based on previous studies that describe the average habitat size for amphibians, ranging from 290 m (Semlitsch & Bodie, 2003) to 500 m (Canessa & Parris, 2013). We consider the central point of the buffer to be the pond or stream (collection points). The polygons for each type of roof were redesigned for the SIRGAS 2000 reference system, Universal Transverse Mercator (UTM) projection, zone 22S and we calculated the areas in km^2^.

### Functional traits of tadpoles

For the analysis of functional diversity, we recorded measurements from one to ten tadpoles of each species. Eighteen morphological traits were measured for each species in the study area. The measured functional traits consider the functional characteristics related to the main aspects of the resource acquisition, use of resources and life-history strategies, as follows: tail muscle height (TMH), tail muscle width (TMW), dorsal fin height (DFH), ventral fin height (VFH), body height (BH), body width (BW), body length (BL), oral disk size (ODS), oral disc position (ODP), number of tooth rows (NTR), eye size (ES), eye distance (IED), eye position (EP), nostril size (NS), internal nostril distance (IND), nostril position (NP), spiracle length (SL), spiracle width (SW), spiracle position (SP). We also recorded the position in the water column (benthic, nektonic and suspenser-rasper), the presence of scourge (present/absent), and ontogenetic development stage (Figure 2; Table 2). The classification of the tadpole position in the water column was defined according to the literature.

**FIGURE 2.**
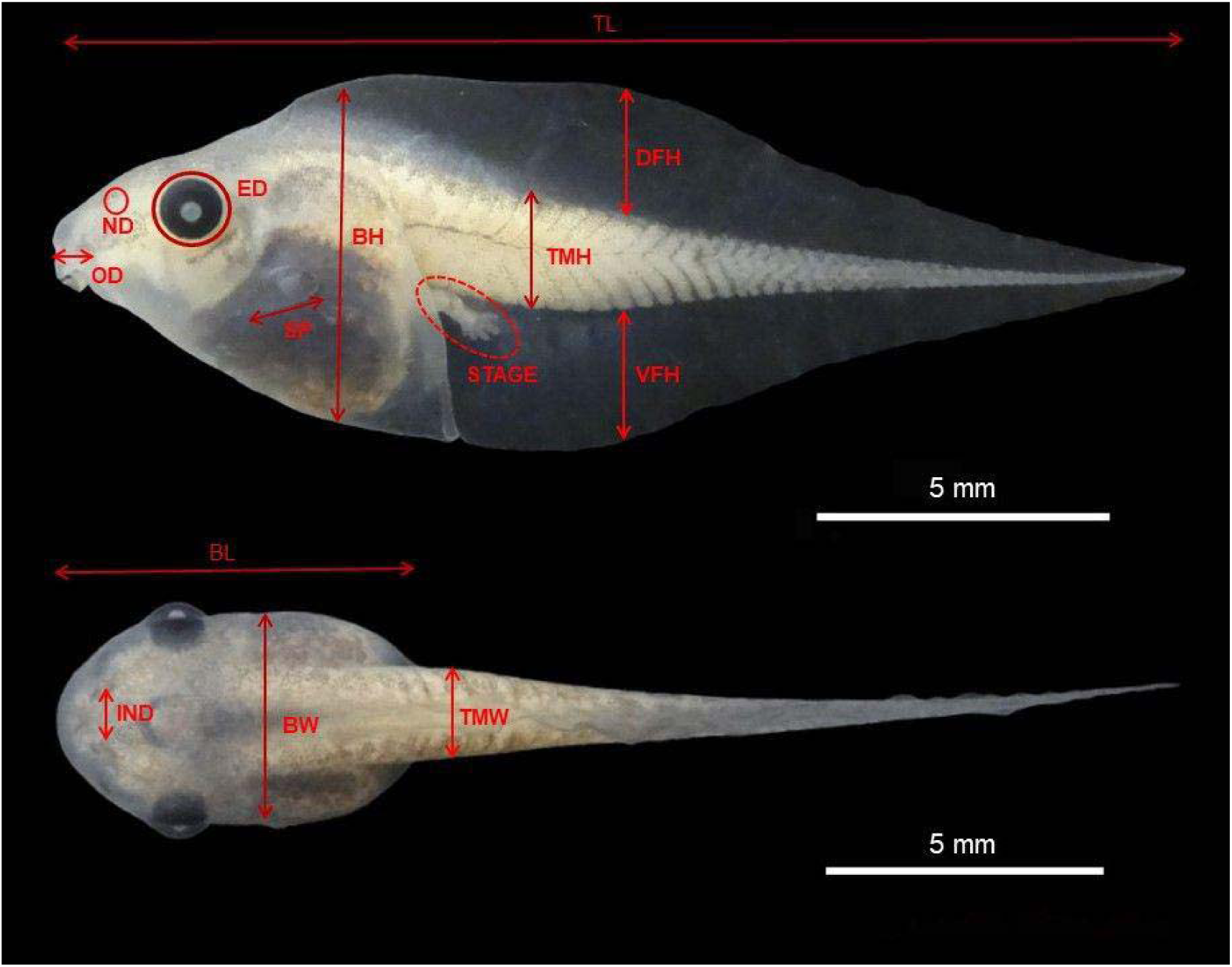
Morphological metrics evaluated in anuran tadpoles: total length (TL); body length (BL); tail muscle height (TMH); tail muscle width (TMW), dorsal fin height (DFH), ventral fin height (VFH), body height (BH), body width (BW), oral disk size (ODS), oral disc position (ODP), number of tooth rows (NTR), eye size (ES), eye distance (IED), eye position (EP), nostril size (NS), internal nostril distance (IND), nostril position (NP), spiracle length (SL), spiracle width (SW), spiracle position (SP). In this picture: Dorsal view and lateral view of *Scinax fuscovarius* tadpoles. (Photographed by Brena da Silva Gonçalves).

**TABLE 2.**
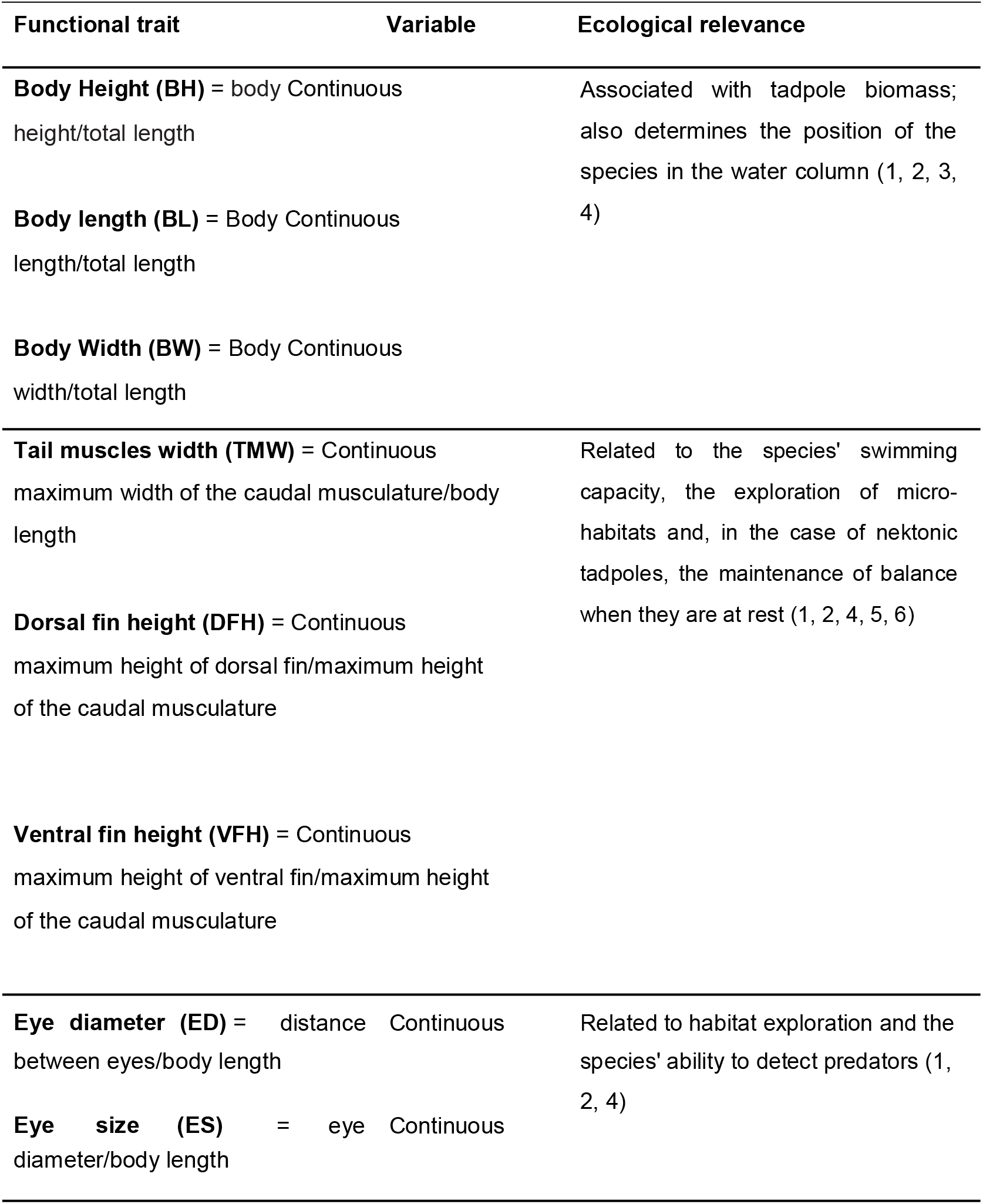
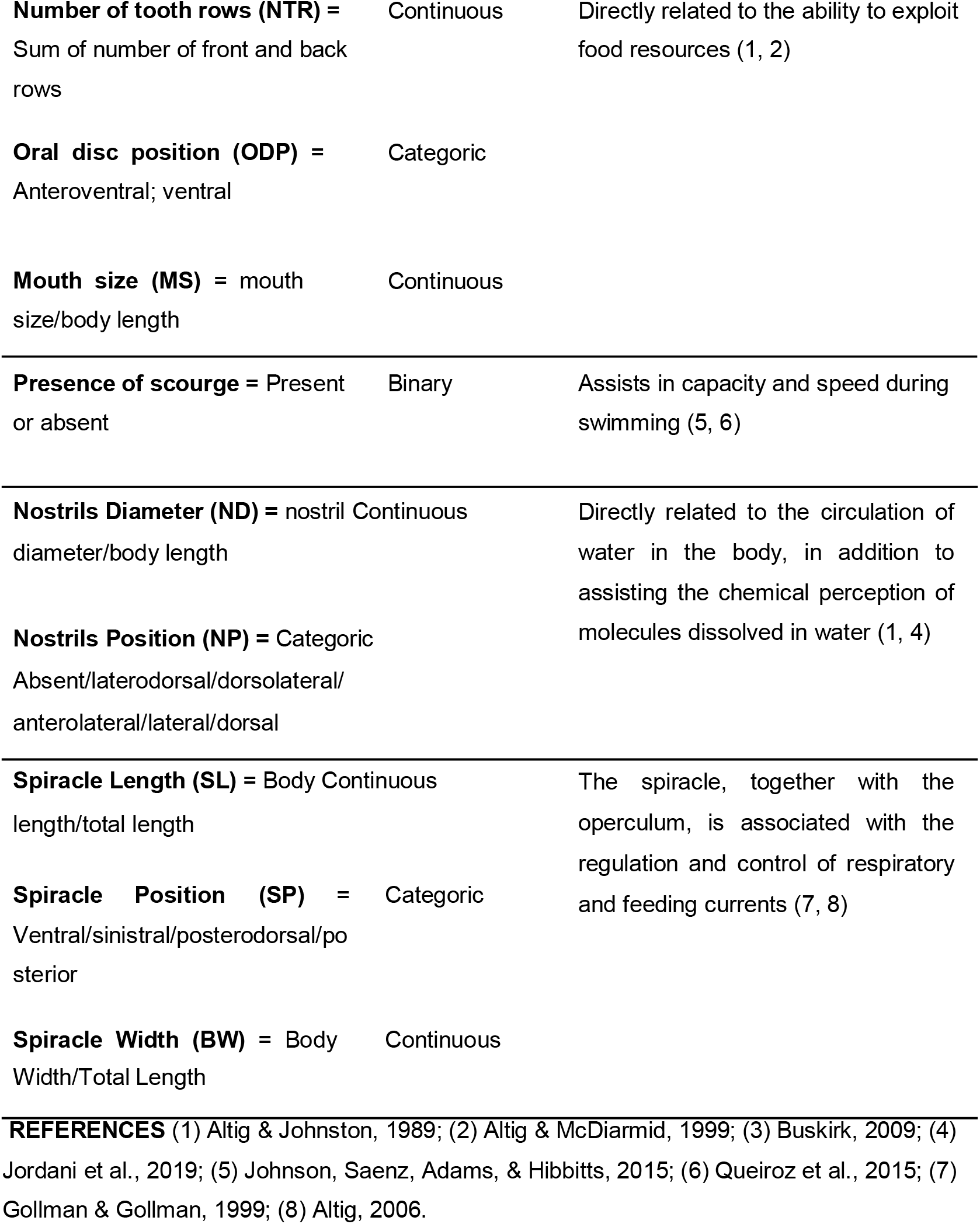
Functional traits measured from different functional characteristics of tadpoles present in the sampled waterbodies.

## DATA ANALYSIS

Before carrying out the statistical analyses, we transformed our quantitative datasets (functional traits and environmental variables) through a natural log transformation; this procedure allows obtaining a normal distribution of the different datasets. Subsequently, we used a combination of the RLQ and the fourth corner methods for assessing the responses of the set of tadpole traits to environmental variation (Dolédec, Chessel, Ter Braak, & Champely, 1996; Dray, Choler, Dolédec, Peres-Neto, Thuiller, Pavoine, & Ter Braak, 2016). In a general view, the proposed RLQ approach is an analysis that performs ordering analyses based on the combination of the following data matrices: an environmental matrix (R; site x environment), a species-by-sites matrix (L; site x species), and a functional-trait distance matrix (Q; species x trait). Matrices R and Q are linked by matrix L.

Before the extended RLQ analysis, we ran the extended version of the fourth-corner approach with 9999 permutations to test the correlations between functional traits and environmental variables. For this, we applied the null model 6 (which fixes the level of type I error; Dray et al., 2016). To prepare the matrices for the extended RLQ analysis, all matrices were analyzed separately with different ordinations: the species-by-site matrix (L) was analyzed using a canonical analysis; the environmental matrix (R) was analyzed by principal component analysis (PCA); finally, the trait distance matrix (Q) and phylogenetic distance matrix P were analyzed by principal coordinate analysis (PCoA). The RLQ analysis and the fourth-corner test were implemented using the packages “spdep” (Bivand, Pebesma, & Gómez-Rubio, 2008) and “ade4” (Dray & Dufour, 2007) of R software.

We run the statistical procedures described above separately for each set of environmental data (i.e., one analysis for local environmental data, and another for landscape data) and for each type of aquatic system (i.e., ponds and streams). To avoid the inclusion of non-significant environmental variables in our models, we performed the analytical procedures described above in two steps. In the first step, we run the analyses containing the full model of environmental variables for each dataset (that is, local and landscape descriptors). Subsequently, we performed a new analysis and included only the environmental variables that showed significant relationships with the functional traits of the tadpoles (i.e., a selected model). The percentage of co-inertia is available as the link between functional traits and environmental descriptors.

## RESULTS

### Community composition

We recorded a total of 22 species, including one non-native, the American Bullfrog (*Lithobates catesbeianus*), with 14 species found in ponds and 19 in streams. The species belong to eight families and 10 genera. Hylidae was the most representative family, with four genera (*Aplastodiscus, Boana, Dendropsophus* and *Scinax*). The genera *Elachistocleis, Proceratophrys* and *Lithobates* occurred only in ponds and the genus *Crossodactylus* only in streams (see Supporting Information Table S1).

### Patterns of trait-environment relationships in ponds

The percentage of co-inertia explained by the two first axes of the fitted RLQ was 75% for the model with local environmental descriptors and 80% for the model with landscape descriptors (Table 3). However, only the model with local environmental descriptors was significant (Std. observed = 3.69, p = 0.001; see Supporting Information Table S2).

**TABLE 3.**
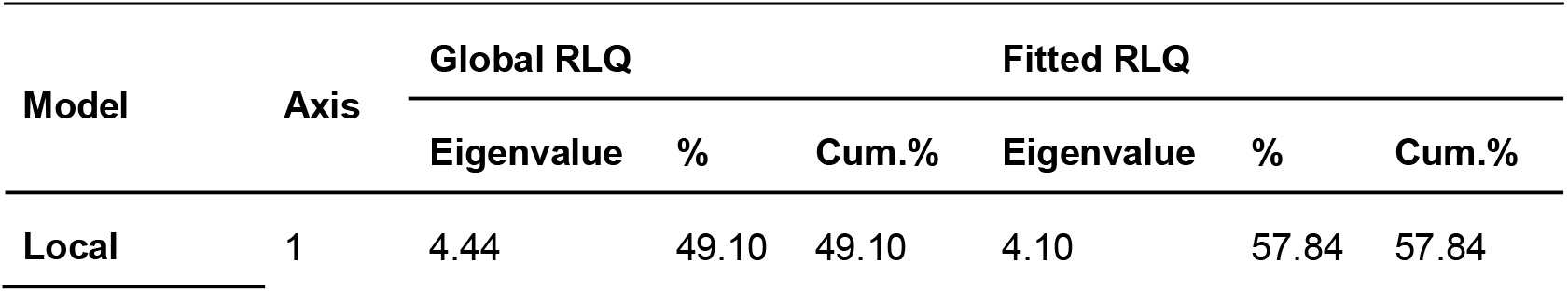
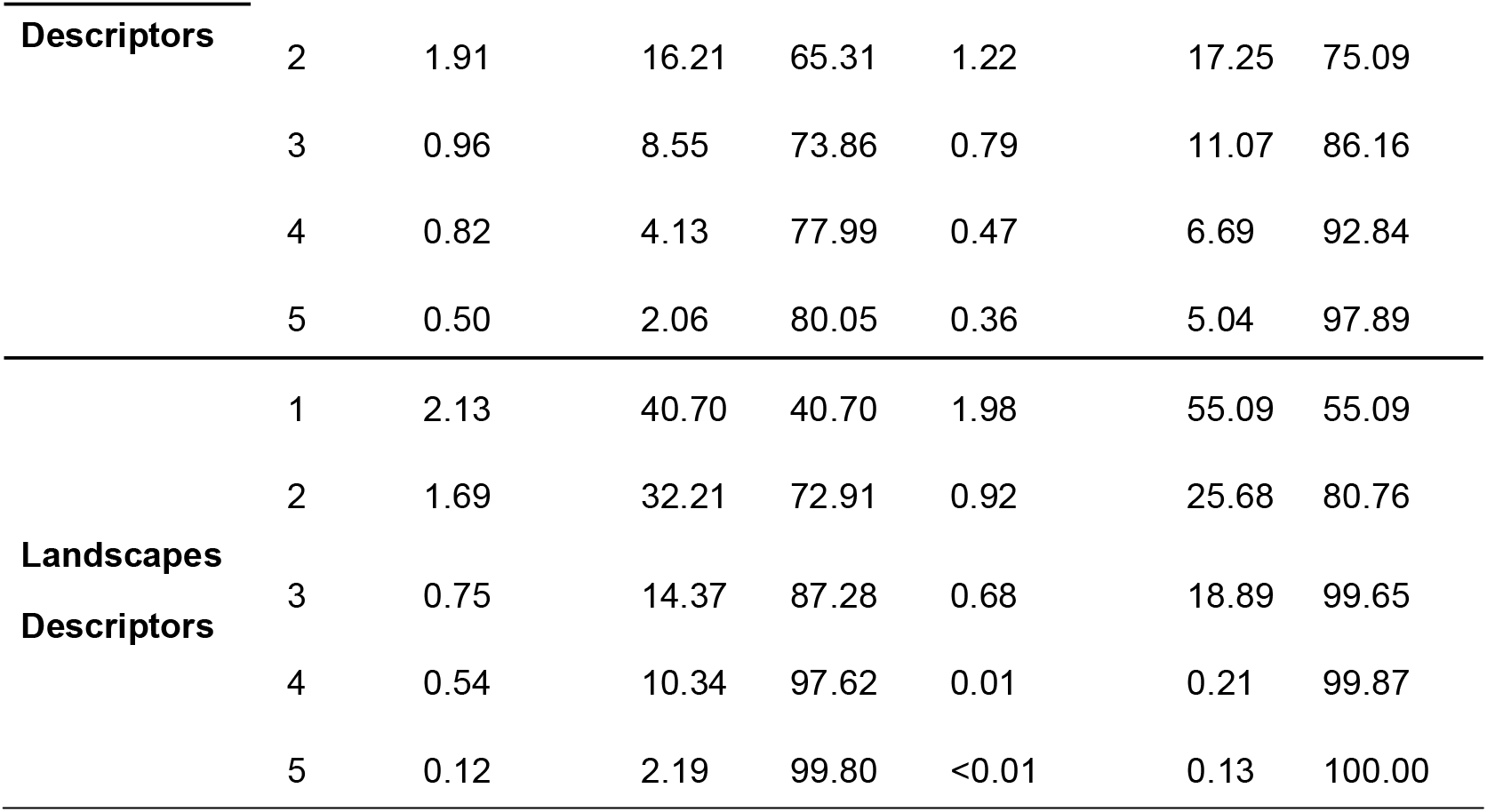
RLQ results from ponds, model of local descriptors, and landscapes descriptors at 28 waterbodies in areas of Atlantic Forest in southern Brazil.

Figure 3a-d presents the patterns of trait-environment relationships observed in ponds. The first RLQ axis had the strongest correlation with the local environmental descriptors pond depth (−2.570; p = 0.006) and water pH (−2.346; p = 0.008), and with the functional traits related to the body (SP POSTERODORSAL: −2.898, p = 0.01), eyes (ED: −1.981, p = 0.04; EP DORSAL: −1.416, p = 0.04), nostrils (NP DORSAL: −1.531, p = 0.02) and tail (DFH: 2.224, p = 0.02; Fl absent: −2.732, p = 0.01). The second RLQ axis had the strongest correlation with trees (−1.397; p = 0.05), water conductivity (−1.650; p = 0.05) and with the functional traits related to nostrils (NP absent: −0.996, p = 0.03) and tails (DFH: 2.224, p = 0.02; see Supporting Information Tables S3 and S4). Benthic tadpoles were more associated with high values of water pH, while nektonic tadpoles were more associated with the presence of trees, although no significant relation of this local environmental descriptor was detected for any of the measured functional traits. Finally, deeper water was more associated with the suspension-rasper tadpoles (Figure 3 a-b). The significance tests of trait-environment relationships are presented in Figure 4a.

**FIGURE 3.**
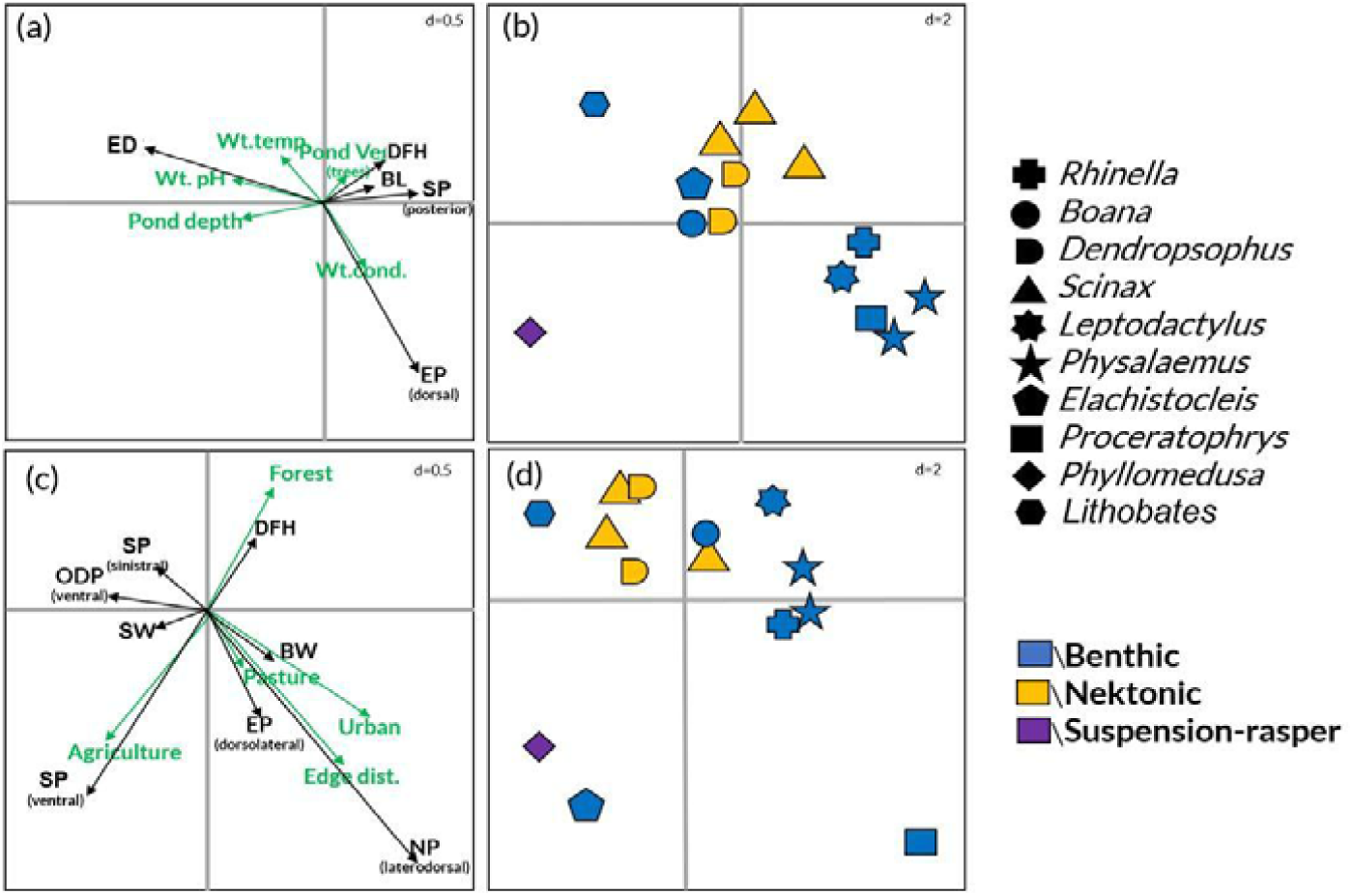
Ordination of tadpoles’ functional traits and ponds’ local descriptors (a), morphological traits and local descriptors (b), ordination of functional traits and landscapes descriptors (c) anuran genera and tadpoles’ trophic guild (d) result of the RLQ analysis. Genera (b and d) are presented by symbols and tadpole trophic guilds by colors: blue = benthic; yellow = nektonic; purple = suspension-rasper. In (a) ED = Eye diameter; EP (dorsal) = Eye position dorsal; DFH = Dorsal fin height; BL= Body length; SP (posterior) = Spiracle position posterior; Wt.temp.= Water temperature; Wt.pH = Water pH; Wt.cond.= Water conductivity; Pond.veg.(grass) = Pond vegetation grass. In (c) EP (lateral) = Eye position lateral; SP (ventral)= Spiracle position ventral; SP (sinistral)= Spiracle position sinistral; NP (dorsolateral) = Nostril position dorsolateral; NP (laterodorsal) = Nostril position laterodorsal; ODP (ventral) = Oral disc position (ventral); SW= spiracle width; DFH = Dorsal fin height; Edge dist.= Edge distance.

**FIGURE 4.**
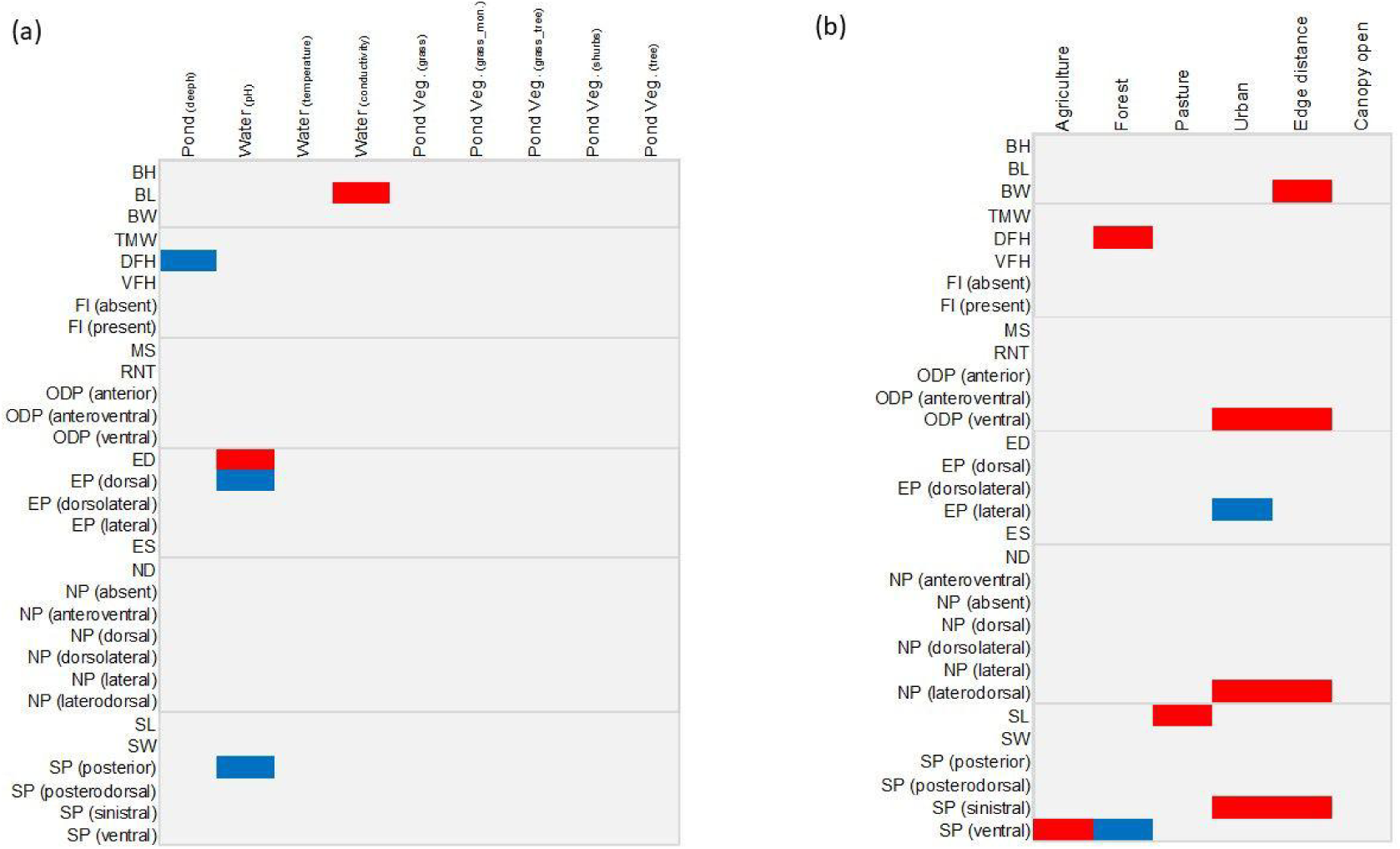
Schemes representing (a) the associations between the tadpoles’ functional traits and ponds’ local descriptors and (b) the associations between the tadpoles’ functional traits and ponds’ landscapes descriptors. Colorless cells represent non-significant associations. Positive and negative associations are represented in blue and red, respectively. The lines show the attribute categories of the body, tail, mouth, eyes, nostrils and spiracle. For functional traits, see Table 2. Abbreviation of morphological traits: BH = Body height; BL= Body length; BW = Body Width; ED = Eye diameter; EP (dorsal) = Eye position dorsal; EP (dorsolateral) = Eye position dorsolateral; EP (lateral) = Eye position lateral; ES = Eye size; FL (absent) = Scourge absent; FL (present) = Scourge present; MS = Mouth size; RNT = Number of tooth rows; ND = Nostril diameter; NP (absent) = Nostril position absent; NP (anterodorsal) = Nostril position anteroventral; NP (dorsal) = Nostril position dorsal; NP (dorsalateral) = Nostril position dorsalateral; NP (lateral) = Nostril position lateral; NP (laterodorsal) = Nostril position laterodorsal; ODP (anterior) = Oral disc position anterior; ODP (anteroventral) = Oral disc position anteroventral; ODP (ventral) = Oral disc position ventral; SL = Spiracle length; SW = Spiracle width; SP (posterior) = Spiracle position posterior; SP (posterodorsal) = Spiracle position posterodorsal; SP (sinistral) = Spiracle position sinistral; SP (ventral) = Spiracle position ventral; TMW = Tail muscle width; DFH = Dorsal fin height; VFH = Ventral fin height. Acronyms of environment attributes in (a): Pond (deeph) = pond water depth; Wt (pH) = Water pH; Wt (temp) = Water temperature; Wt (cond)= Water conductivity; Pond Veg. (grass) = Pond vegetation grasses; Pond Veg. (grass.mon) = Pond vegetation grass monoculture; Pond Veg. (grass.tree) = Pond vegetation grasses and trees; Pond Veg. (shrubs) = Pond vegetation shrubs; Pond Veg. (tree) = Pond vegetation tree.

The first two axes of the RLQ model for the landscape descriptors of ponds accounted for, respectively, 55% and 26% of the variance. Despite this, this model was not statistically significant (Std. observed = 1.447, p = 0.07; Table 3, Figure 3 c-d). However, the fourth-corner test showed significant relationships between some landscape descriptors and functional traits (Figure 4b). We observed a negative relationship between forest and dorsal fin height (DFH); agriculture and spiracle position (SP ventral); pasture and spiracle width (SW); urban and oral disc position (ODP ventral), spiracle position (SP sinistral), and nostril position (NP laterodorsal); edge distance and body width (BW), oral disc position (ODP ventral), spiracle position (SP sinistral) and nostril position (NP laterodorsal). Finally, we observed positive relationships between forest and spiracle position (SP ventral), and urban and eye position (EP lateral) (Supporting Information Table S5).

### Patterns of trait-environment relationships in streams

The percentage of co-inertia explained by the two first axes of the fitted RLQ was 73% for the model with local descriptors, and 78% for the model with landscape descriptors (Table 4). Similar to the patterns observed in ponds, only the model with local descriptors was significant (Std. observed = 3.54, p = 0.001; see Supporting Information Table S6).

**TABLE 4.** RLQ results from streams, model of local descriptors, and landscapes descriptors at 28 waterbodies in areas of Atlantic Forest in southern Brazil.

| **Model** | **Axis** | **Global RLQ** |  |  | **Fitted RLQ** |  |  |
| --- | --- | --- | --- | --- | --- | --- | --- |
|  |  | Eigenvalue | % | Cum.% | Eigenvalue | % | Cum.% |
| **Local**<br>**Descriptors** | 1 | 9.50 | 46.15 | 46.15 | 8.40 | 48.46 | 48.46 |
|  | 2 | 5.56 | 27.04 | 73.19 | 4.25 | 24.50 | 72.96 |
|  | 3 | 2.31 | 11.20 | 84.39 | 2.03 | 11.73 | 84.69 |
|  | 4 | 1.23 | 5.99 | 90.39 | 0.97 | 5.58 | 90.27 |
|  | 5 | 0.98 | 4.78 | 95.17 | 0.85 | 4.92 | 95.19 |
| **Landscapes**<br>**Descriptors** | 1 | 4.72 | 48.43 | 48.43 | 2.51 | 56.84 | 56.84 |
|  | 2 | 2.27 | 23.30 | 71.73 | 0.96 | 21.67 | 78.51 |
|  | 3 | 1.11 | 11.43 | 83.16 | 0.46 | 10.37 | 88.88 |
|  | 4 | 0.67 | 6.93 | 90.09 | 0.35 | 7.81 | 96.69 |
|  | 5 | 0.47 | 4.79 | 94.89 | 0.12 | 2.71 | 99.40 |

Figure 5a-d presents the patterns of trait-environment relationships observed in streams. The first RLQ axis had the strongest correlation with the local environmental descriptors water pH (−2.453, p = 0.008), water temperature (−2.397, p = 0.01), water conductivity (2.162, p = 0.01), total alkalinity (2.162, p = 0.03), Alk HCO3 (2.122, p = 0.03) and stream mud+rocks (2.162, p = 0.01; Supporting Information Table S7), and with the functional traits related to the body (BW: −2.258, p = 0.02; SP posterodorsal: − 2.444, p = 0.01), eyes (EP dorsal: −1.487, p = 0.05; EP lateral: −1.529, p = 0.03), mouth (NTR: −2.123, p = 0.02; ODP anteroventral: −2.325, p = 0.03) and tail (Fl present: −1.726, p = 0.02; VFH: 2.045, p = 0.04; Supporting Information Table S8). The second RLQ axis had the strongest correlation with the types of stream vegetation (presence of trees: −2.112; p = 0.03; presence of grass+rocks: −1.087; p = 0.04), and with the functional traits related to the mouth (MS: −1.900, p = 0.04). Most benthic tadpoles were more associated with the physicochemical descriptors of water (pH, temperature) and with trees, while the second group of benthic species (consisting of the genera *Aplastodiscus, Boana* and *Lithobathes*) were more associated with the physicochemical descriptors of water NO3, Alk HCO3, total alkalinity and conductivity. Nektonic tadpoles were more associated with stream area and water transparency. Finally, the suspension-rasper tadpoles were more associated with the presence of aquatic vegetation on the stream substrate (Figure 5a-b). The significance tests of trait-environment relationships are presented in Figure 6a.

**FIGURE 5.**
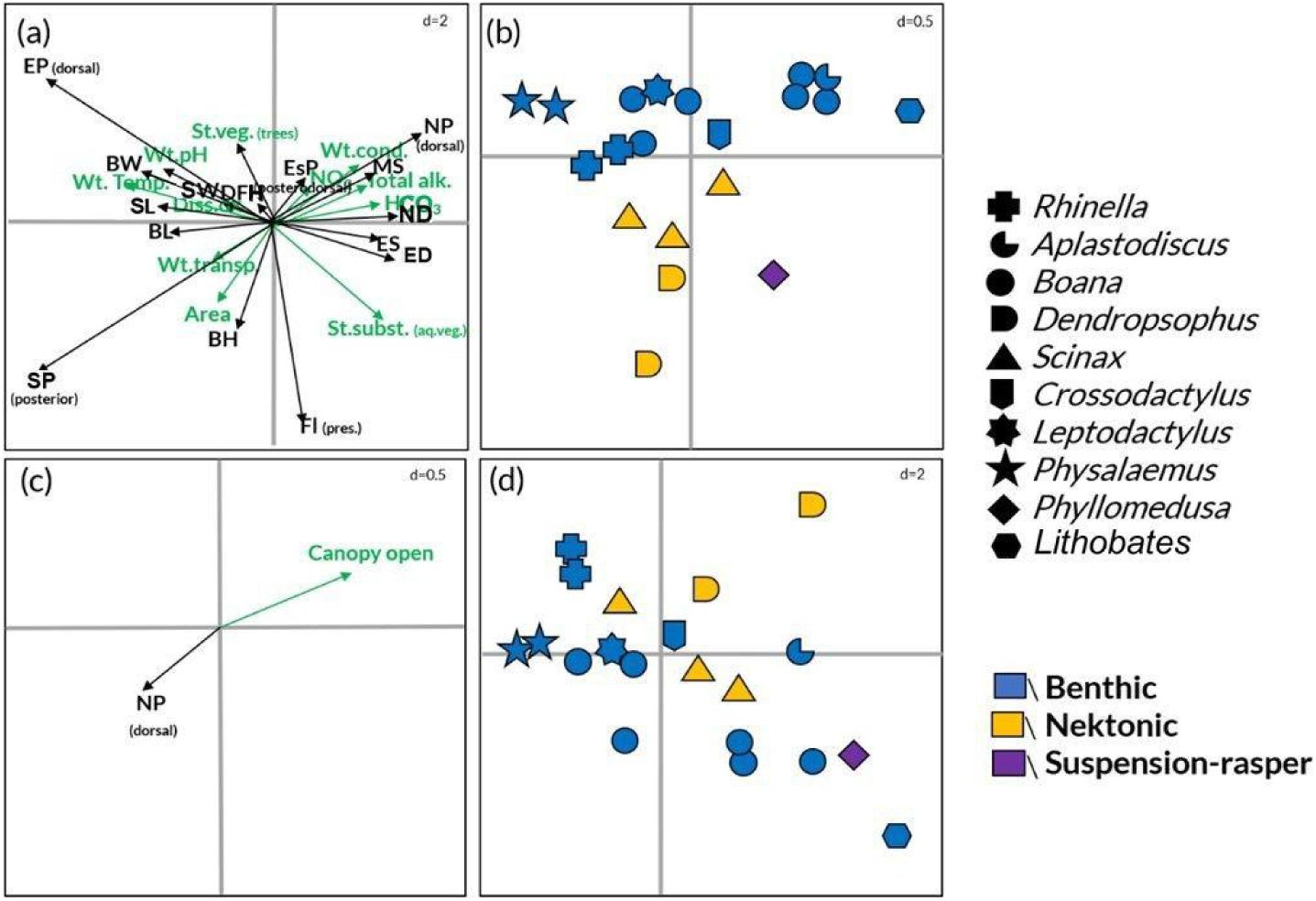
a-b (local descriptors), c-d (landscape descriptors). Ordination of functional traits and local descriptors (a), anuran genera and habitat use (b), ordination of functional traits and landscapes descriptors (c), anuran genera and habitat use (d) result of the RLQ analysis. Genera (b and d) are represented by symbols and tadpole trophic guilds by colors: blue = benthic; yellow = nektonic; purple = suspension-rasper. In (a) ED = Eye diameter; EP (dorsal) = Eye position dorsal; DFH = Dorsal fin height; BL= Body length; SP (posterior) = Spiracle position posterior; Wt.temp.= Water temperature; Wt.pH = Water pH; Wt.cond.= Water conductivity; Pond.veg.(grass)= Pond vegetation grass. In (c): EP (lateral) = Eye position lateral; SP (ventral)= Spiracle position ventral; SP (sinistral) = Spiracle position sinistral; NP (dorsolateral)= Nostril position dorsolateral; NP (laterodorsal) = Nostril position laterodorsal; ODP (ventral) = Oral disc position ventral; SW= spiracle width; DFH = Dorsal fin height; Edge dist.= Edge distance.

**FIGURE 6.**
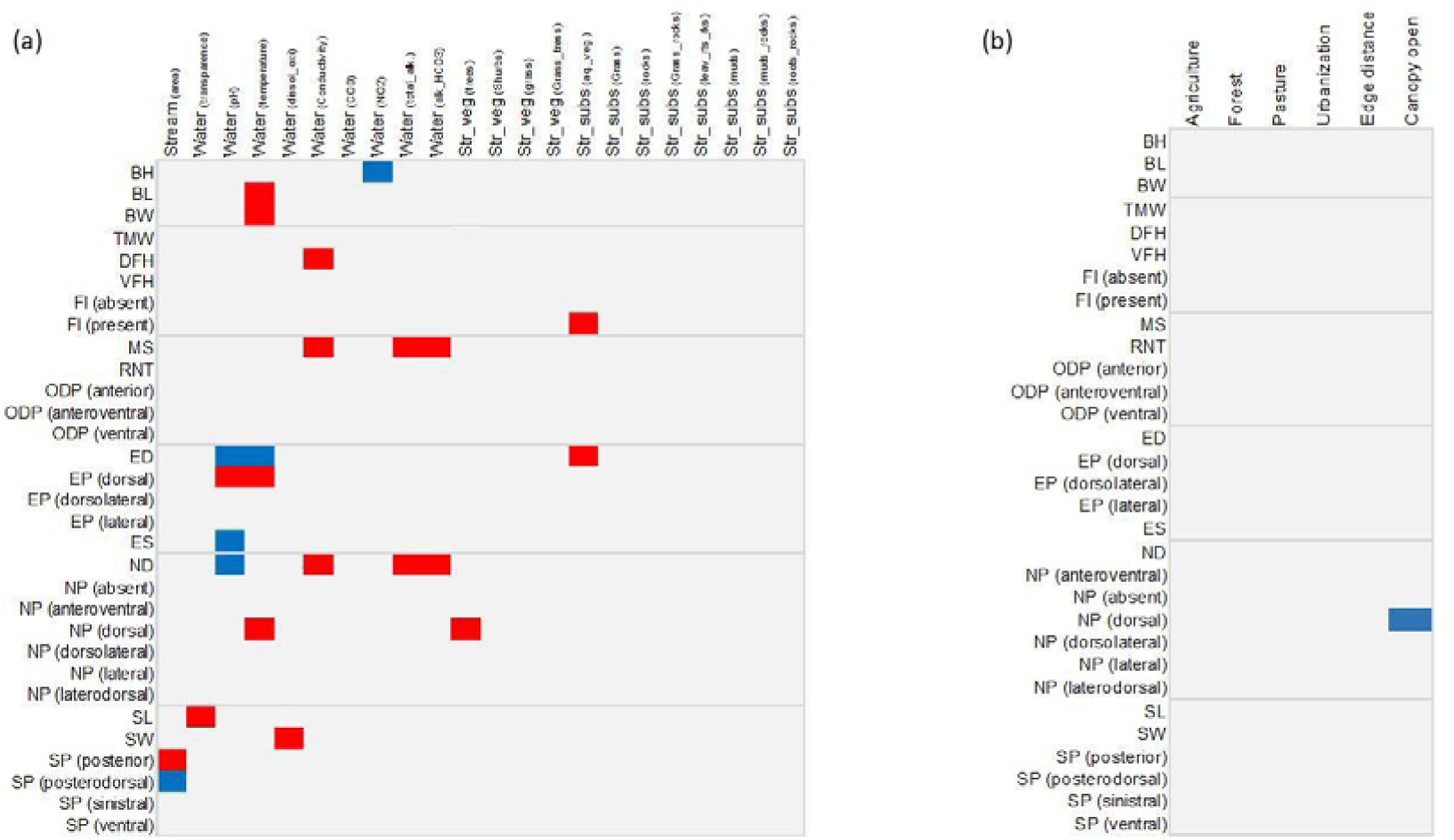
Schemes representing (a) the associations between the tadpoles functional traits and stream local descriptors. (b) the associations between the tadpoles functional traits and stream landscapes descriptors. Colorless cells represent non-significant associations. Positive and negative associations are represented in blue and red, respectively. The lines show the attribute categories of the body, tail, mouth, eyes, nostrils and spiracle. For functional traits, see Table 2. Abbreviations of morphological traits: BH = Body height; BL= Body length; BW = Body Width; ED = Eye diameter; EP (dorsal) = Eye position dorsal; EP (dorsolateral) = Eye position dorsolateral; EP (lateral) = Eye position lateral; ES = Eye size; FL (absent) = Scourge absent; FL (present) = Scourge present; MS = Mouth size; RNT = Number of tooth rows; ND = Nostril diameter; NP (absent) = Nostril position absent; NP (anterodorsal) = Nostril position anteroventral; NP (dorsal) = Nostril position dorsal; NP (dorsalateral) = Nostril position dorsalateral; NP (lateral) = Nostril position lateral; NP (laterodorsal) = Nostril position laterodorsal; ODP (anterior) = Oral disc position anterior; ODP (anteroventral) = Oral disc position anteroventral; ODP (ventral) = Oral disc position ventral; SL = Spiracle length; SW = Spiracle Width; SP (posterior) = Spiracle position posterior; SP (posterodorsal) = Spiracle position posterodorsal; SP (sinistral) = Spiracle position sinistral; SP (ventral) = Spiracle position ventral; TMW = Tail muscle width; DFH = Dorsal fin height; VFH = Ventral fin height. Abbreviations of environment attributes, in (a): Stream (area) = Stream area; Wt (tranparency) = Water transparency; Wt (pH) = Water pH; Wt (temperature) = Water temperature Wt (dissol.oxi)= Water dissolved oxygen; Wt (cond)= Water conductivity; Wt (COD)= Water chemical oxygen demand; Wt (NO_2_)= Water nitrite; Wt (Total_alk) = Water total alkalinity; Wt (Alk_HCO_3_) = Bicarbonate alkalinity; Str_Veg (trees) = Stream vegetation trees; Str_Veg (shrubs) = Stream vegetation shrubs; Str_Veg (grass) = Stream vegetation grasses; Str_Veg (grass_trees) = Stream vegetation grasses and trees; Str_subs (Aq._veg) = Stream substrate with aquatic vegetation; Str_subs (grass) = Stream substrate with grasses; Str_subs (rocks) = Stream substrate with rocks; Str_subs (grass_rocks) = Stream substrate with grasses and rocks; Str_subs (leav_rts_rks) = Stream substrate with leaves, roots and rocks; Str_subs (muds) = Stream substrate with muds; Str_subs (muds_rocks) = Stream substrate with muds and rocks; Str_subs (roots_rocks) = Stream substrate with roots and rocks.

The two first axes of the RLQ model to the landscape descriptors of streams accounted for, respectively, 57% and 22% of the variance. Despite this, this model was not statistically significant (Std. observed = 1.33, p = 0.09; Table 4; Supporting Information Tables S9 and S10). However, the fourth-corner test showed significant relationships between canopy opening and the functional trait nostril position dorsal (NP dorsal) (Figure 6b; Supporting Information Table S11).

## Discussion

Our results indicated that the local habitat variables are more relevant than the landscape variables to define functional traits in the communities of both ponds and streams. Both communities are expected to be exposed to different selective pressures due to differences between ponds and streams regarding abiotic (Smith, Renwick, Bartley, & Buddemeier, 2002; Fairchild & Velinsky, 2006; Hoeinghaus, Winemiller, & Birnbaum, 2007) and biotic components (Schriever & Lytle, 2016; Jordani et al., 2017). However, we observed differences even in the type of association that each environmental descriptor caused on the functional trait of tadpoles between these two waterbodies types. It is interesting to notice that, despite the evident differences in terms of functional traits, the species composition in these two systems was similar. The exclusivity of some species had a strong relationship with reproductive mode, i.e., species that are exclusive to stream communities lay their eggs only in this type of habitat (*Boana curupi* and *Crossodactylus schmidti*; Carrizo, 1991; Caldart, Iop, & Cechin, 2014), with the same can be said for species that occur mainly in ponds (*Elachistocleis, Proceratophrys* and *Lithobates*) (Rodrigues, Lopes & Uetanabaro, 2003; Both, 2012).

In ponds, water depth was an environmental variable associated with a large number of functional traits. It was, for example, positively associated with the dorsal height of the tail fin, which would affect the swimming of tadpoles. Since fin height contributes to displacement in the water column (Johnson et al., 2015; Jordani et al., 2019), this would be an important factor, especially for nektonic species (e.g., *Dendropsophus* and *Scinax*), as it increases their ability to exploit different resources in the water column (Marques & Nomura, 2018). The relationship between depth and functional traits deserves attention since the functional diversity of tadpoles tends to increase in medium depths, suggesting that it is surrounded by a complex set of ecological interactions (Queiroz et al., 2015). However, attributing functional traits to a single environmental factor means approaching the ecological and adaptive processes that may be acting on this process in a simplified manner. Alternatively, we can associate pond depth to the pond’s cycle. In the case of shallower ponds, depth reflects the pond’s longevity (hydroperiod), which defines the possibility of a community occupying a place (Jordani et al., 2017; Meyer, Franklin, & Cramp, 2020). In this case, the reduction in depth would be an indicator of a pond’s longevity, giving the tadpoles environmental clues for the timing of their metamorphosis. The speed at which temporary ponds dry out interferes with the speed that is needed for metamorphosis and in the size of the newly metamorphosed (Wellborn et al., 1996; Babbitt, Baber, & Tarr, 2003; Johnson et al., 2015). The fact that depth was an important component only in ponds may be related to the low stability and predictability of the ponds that were sampled in our study area. Hydroperiod also affects the processes of predation and/or competition (Simpkins, Shuker, Lollback, Castley, & Hero, 2013; Wellborn et al., 1996; Werner et al., 2007; Melo et al., 2018). This can justify the association that we recorded between water depth and morphology of the eyes (predator detection), tail (mobility, escape velocity) and body size (susceptibility to predation).

However, associating morphology with specific environmental factors is a complex and risky task (Lopes, Serra, Piorski, & De Andrade, 2020). Nevertheless, some generalizations can be made. The position of the eyes, for example, would be related to habitat evaluation, height of the tail’s dorsal fin and filament, and locomotion capacity (Altig & Johnston, 1989; Altig & McDiarmid, 1999, Johnson et al., 2015; Queiroz et al., 2015; Jordani et al., 2019), while nostrils would be related to physiological activities of osmoregulation (Altig & Johnston, 1989; Jordani et al., 2019), and the morphology of the mouth apparatus would be associated with feeding (Altig & Johnston, 1989; Altig & McDiarmid, 1999). Thus, it is possible to expect morphological changes to result from processes of physiological compensations to a certain environmental stressor. We could, therefore, speculate about the morphology-physiology relationship and changes in functional attributes. There are indications, for example, that an increase in tail height may be a response to predation pressure (McCollum & Leimberger, 1997; Relyea, 2003; Relyea & Hoverman, 2003), and the effects of the predation risk on microhabitat selection are well known in aquatic organisms. Vegetation, for example, enhances the use of microhabitats with greater refuge availability (Diaz-Paniagua, 1987; Kopp, Wachlevski, & Eterovick, 2006), interfering in the predation process. Other evaluated local variables, such as pH, water conductivity and presence of arboreal vegetation were associated with functional traits of the nostrils, spiracle, eyes and tail morphology. Physicochemical components of water may affect the physiology and development of tadpoles, which reflects in compensation mechanisms. Low pH values may affect tadpole growth (Pierce, 1985; Farquharson et al., 2016; Meyer, Franklin, & Cramp, 2020). The traits of nostrils and spiracle are related to the tadpoles’ respiratory and regulatory physiology, water intake and flow into the body (Gollman & Gollman, 1999; Altig, 2006), which has a strong connection the physicochemical properties of water. Water conductivity, in turn, is related to the susceptibility to diseases (e.g., bacteria and the fungus *Batrachochytrium denbrobatidis*; Carey, 1993; Klaver, Peterson, & Patla, 2013).

It is interesting to notice that the physical and chemical descriptors of water revealed to be more relevant for stream communities than for pond communities. In streams, functional traits were associated with a larger set of descriptors, such as pH, temperature, conductivity, total alkalinity, Alk and HCO3. Besides the water descriptors, the most relevant components in streams were linked to the substrate and the characteristics of the margin, such as vegetation type (tree). While the water parameters are more connected to physiological processes, the microhabitat elements may be more related to foraging and shelter provision. This hypothesis is reinforced by the fact that this set of descriptors showed a strong association with functional traits that could be related to predation susceptibility (body size and morphology of fins and the tail), as well as to habitat exploration and acquisition of food resources (position of the eyes, mouth and oral disc).

Another interesting point is that we found species groups linked to their guilds, but the descriptors that are more associated with them are habitat-specific. In streams, most benthic species are more associated with descriptors of water (pH, temperature, NO3, HCO3, total alkalinity and conductivity) and with trees. On the other hand, in ponds, benthic species were only associated with pH. In streams, nektonic species were more associated with stream area and greater water transparency, while in ponds they were associated only with the presence of trees. Suspension-rasper tadpoles, in turn, were more associated with the presence of aquatic vegetation on the stream substrate in streams and with depth in ponds.

Although we cannot associate these differences with elements of their life history, the fact that there is a relationship between guilds and different habitat parameters and respective functional traits is extremely relevant. Some derivations can be listed, although speculatively. Temperature, for example, is one of the main local variables that influence the physical, chemical and biological processes of streams (Caissie, 2006). Studies relating tadpole growth to temperature show that the latter affects tadpole development (see Browne & Edwards, 2003), with growth speeding up under ideal temperatures for tadpole growth (Maciel & Juncá, 2009). Because ponds are lentic systems, they are expected to be more stable environments regarding water parameters than streams. Depth would be a variable defining the stability level of this system: the deeper, the more stable. However, the surrounding environment of ponds and streams has a great potential to change the physical and chemical characteristics of water (Sipaúba-Tavares et al., 2007; Mansano et al., 2012, 2014). The vegetation at the margins of streams and ponds acts as an important resource and filter of organic matter for aquatic environments (Lecerf, Dobson, Dang, & Chauvet, 2005), affecting energy flow and the whole local trophic web (Antoniazzi, López, Lorenzón, Saigo, Devercelli, & et al., 2020). Our results indicated possible adjustments in the functional traits related to physicochemical characteristics of the water and microhabitat used by tadpoles in the waterbodies. These adjustments, highlighted by differences in functional traits between ponds and streams, indicate ecological adjustments in the face of different environmental conditions. These adjustments can be seen as an important survival tool in systems that are susceptible to fast changes in their conditions. Rain, sediment input, arrival of predators are important elements in the stability of the habitat of tadpoles. Since tadpoles have a short time to complete their metamorphosis, the possibility of quick responses in the face of dynamic and somewhat unpredictable environmental conditions would be an important tool for the reproductive success of anurans.

## ACKNOWLEDGEMENTS

We are thankful to all private owners that allowed us to access their properties and the state environmental agencies that authorized the research in the conservation units. We also thank the Coordenação de Aperfeiçoamento de Pessoal de Nível Superior (CAPES) - Finance Code 001, via a Master’s degree fellowship to DB, and the Pe. Theobaldo Frantz Fund via a doctorate fellowship to RCS. Piter Kehoma Boll helped with the English translation and editing of the manuscript.

