## Supplementary files for "Trait-environment relationship in tadpoles of the southern Atlantic Forest"

SUPORTING INFORMATION

Table S1: TadpoleT

**Suporting Information**

Table S1: Tadpole abundance for 28 waterbodies in areas of Atlantic Forest in southern Brazil, recorded from October 2018 to March 2019. P = Pond; S = Stream

| Family/Species | 1 | 2 | 3 | 4 | 5 | 6 | 7 | 8 | 9 | 10 | 11 | 12 | 13 | 14 | 15 | 16 | 17 | 18 | 19 | 20 | 21 | 22 | 23 | 24 | 25 | 26 | 27 | 28 |
| --- | --- | --- | --- | --- | --- | --- | --- | --- | --- | --- | --- | --- | --- | --- | --- | --- | --- | --- | --- | --- | --- | --- | --- | --- | --- | --- | --- | --- |
|  | P S S P S P S P S P S P S P S S P S P S S P S S P S P | | | | | | | | | | | | | | | | | | | | | | | | | | | S |
| BUFONIDAE |  |  |  |  |  |  |  |  |  |  |  |  |  |  |  |  |  |  |  |  |  |  |  |  |  |  |  |  |
| *Rhinella henseli* (Lutz, 1934) | 0 | 0 | 0 | 0 | 119 | 0 | 0 | 0 | 0 | 0 | 12 | 0 | 0 | 0 | 0 | 0 | 0 | 0 | 0 | 0 | 0 | 0 | 0 | 0 | 0 | 0 | 0 | 0 |
| *Rhinella icterica* (Spix, 1824) | 0 | 0 | 0 | 0 | 0 | 400 | 1 | 0 | 0 | 0 | 0 | 40 | 0 | 0 | 0 | 0 | 0 | 19 | 0 | 0 | 0 | 0 | 0 | 0 | 0 | 0 | 0 | 0 |
| HYLIDAE |  |  |  |  |  |  |  |  |  |  |  |  |  |  |  |  |  |  |  |  |  |  |  |  |  |  |  |  |
| *Aplastodiscus perviridis* Lutz 1950 | 0 | 0 | 0 | 0 | 0 | 0 | 1 | 0 | 0 | 0 | 0 | 0 | 0 | 0 | 0 | 0 | 0 | 0 | 0 | 0 | 0 | 0 | 0 | 0 | 0 | 0 | 0 | 0 |
| *Boana* cf. *curupi* | 0 | 0 | 0 | 0 | 0 | 0 | 0 | 0 | 0 | 0 | 0 | 0 | 0 | 0 | 0 | 0 | 0 | 0 | 0 | 0 | 55 | 0 | 30 | 37 | 0 | 0 | 0 | 0 |
| *Boana curupi* (Garcia, Faivovichi and Haddad, 2007) | 0 | 0 | 0 | 0 | 0 | 0 | 0 | 0 | 0 | 0 | 0 | 0 | 0 | 0 | 50 | 98 | 0 | 0 | 0 | 0 | 43 | 0 | 6 | 41 | 0 | 0 | 0 | 0 |
| *Boana faber* (Wied - Neuwied 1821) | 0 | 0 | 0 | 0 | 0 | 0 | 0 | 11 | 1 | 53 | 1 | 0 | 0 | 124 | 1 | 0 | 14 | 91 | 48 | 0 | 0 | 10 | 0 | 0 | 40 | 6 | 0 | 0 |
| *Boana leptolineata* (Braun and Braun, 1977) | 0 | 0 | 0 | 0 | 0 | 0 | 0 | 0 | 0 | 0 | 0 | 0 | 0 | 0 | 3 | 6 | 0 | 0 | 0 | 0 | 0 | 0 | 0 | 0 | 0 | 0 | 0 | 0 |
| *Boana prasina* (Burmeister, 1856) | 0 | 67 | 16 | 0 | 0 | 0 | 0 | 0 | 0 | 0 | 0 | 0 | 0 | 0 | 0 | 0 | 0 | 0 | 0 | 0 | 0 | 0 | 0 | 0 | 0 | 0 | 0 | 0 |
| *Boana pulchella* (Duméril and Bibron,1841) | 0 | 9 | 4 | 0 | 0 | 0 | 0 | 0 | 0 | 0 | 0 | 0 | 0 | 0 | 0 | 0 | 0 | 0 | 0 | 0 | 0 | 0 | 0 | 0 | 0 | 0 | 0 | 0 |
| *Dendropsophus microps* (Peters, 1872) | 0 | 0 | 0 | 0 | 12 | 0 | 0 | 0 | 0 | 1 | 4 | 0 | 9 | 0 | 0 | 0 | 0 | 0 | 7 | 0 | 0 | 40 | 0 | 0 | 14 | 0 | 0 | 0 |
| *Dendropsophus minutus* (Peters, 1872) | 5 | 0 | 0 | 0 | 0 | 0 | 0 | 21 | 0 | 0 | 0 | 5 | 14 | 18 | 1 | 0 | 0 | 0 | 53 | 2 | 0 | 0 | 0 | 0 | 2 | 0 | 0 | 0 |
| *Scinax fuscovarius* (Lutz, 1925) | 0 | 0 | 0 | 3 | 0 | 0 | 0 | 28 | 0 | 0 | 0 | 0 | 9 | 0 | 0 | 0 | 0 | 0 | 9 | 0 | 0 | 5 | 0 | 0 | 0 | 0 | 0 | 0 |
| *Scinax granulatus* (Peters, 1871) | 0 | 0 | 3 | 0 | 0 | 0 | 0 | 3 | 0 | 0 | 0 | 5 | 9 | 0 | 2 | 0 | 0 | 0 | 0 | 0 | 0 | 0 | 0 | 0 | 0 | 0 | 0 | 0 |
| *Scinax perereca* Pombal, Haddad and Kasahara, 1995 | 0 | 0 | 0 | 8 | 0 | 0 | 0 | 1 | 0 | 0 | 2 | 0 | 0 | 0 | 0 | 0 | 0 | 0 | 0 | 0 | 0 | 9 | 0 | 0 | 0 | 0 | 90 | 67 |
| HYLODIDAE |  |  |  |  |  |  |  |  |  |  |  |  |  |  |  |  |  |  |  |  |  |  |  |  |  |  |  |  |
| *Crossodactylus schmidti* Gallardo, 1961 | 0 | 0 | 0 | 0 | 0 | 0 | 124 | 0 | 55 | 0 | 0 | 0 | 0 | 0 | 0 | 0 | 0 | 0 | 0 | 0 | 16 | 0 | 0 | 0 | 0 | 118 | 0 | 0 |
| LEPTODACTYLIDAE |  |  |  |  |  |  |  |  |  |  |  |  |  |  |  |  |  |  |  |  |  |  |  |  |  |  |  |  |

1

2

3

4

5

6

7

8

9

10

11

12

13

14

15

16

17

18

19

20

21

22

23

24

25

26

27

28

29

30

31

32

33

34

35

36

37

38

39

40

41

42

43

44

45

46

Freshwater Biology	Page 42 of 60

| *Leptodactylus latrans* (Steffen, 1815) | 0 | 0 | 0 | 0 | 0 | 0 | 0 | 0 | 0 | 19 | 0 | 0 | 0 | 0 | 0 | 0 | 0 | 53 | 0 | 0 | 0 | 0 | 0 | 0 | 0 | 0 | 0 | 0 |
| --- | --- | --- | --- | --- | --- | --- | --- | --- | --- | --- | --- | --- | --- | --- | --- | --- | --- | --- | --- | --- | --- | --- | --- | --- | --- | --- | --- | --- |
| *Physalaemus cuvieri* Fitzinger, 1826 | 6 | 0 | 0 | 1 | 39 | 0 | 0 | 0 | 0 | 6 | 0 | 110 | 53 | 0 | 0 | 0 | 0 | 0 | 28 | 64 | 0 | 0 | 0 | 0 | 0 | 0 | 0 | 0 |
| *Physalaemus* cf. *carrizorum* | 0 | 0 | 0 | 0 | 0 | 0 | 0 | 0 | 0 | 0 | 0 | 2 | 11 | 0 | 0 | 0 | 0 | 0 | 0 | 0 | 0 | 8 | 0 | 0 | 0 | 0 | 0 | 0 |
| MICROHYLIDAE |  |  |  |  |  |  |  |  |  |  |  |  |  |  |  |  |  |  |  |  |  |  |  |  |  |  |  |  |
| *Elachistocleis bicolor* (Guérin-Méneville, 1838) | 0 | 0 | 0 | 0 | 0 | 0 | 0 | 2 | 0 | 0 | 0 | 0 | 0 | 0 | 0 | 0 | 0 | 0 | 0 | 0 | 0 | 0 | 0 | 0 | 0 | 0 | 0 | 0 |
| ODONTOPHRYNIDAE |  |  |  |  |  |  |  |  |  |  |  |  |  |  |  |  |  |  |  |  |  |  |  |  |  |  |  |  |
| *Proceratophrys avelinoi* Mercadal de Barrio and Barrio, 1993 | 0 | 0 | 0 | 0 | 0 | 0 | 0 | 0 | 0 | 0 | 0 | 0 | 0 | 0 | 0 | 0 | 7 | 0 | 0 | 0 | 0 | 0 | 0 | 0 | 0 | 0 | 0 | 0 |
| PHYLLOMEDUSIDAE |  |  |  |  |  |  |  |  |  |  |  |  |  |  |  |  |  |  |  |  |  |  |  |  |  |  |  |  |
| *Phyllomedusa tetraploidea* Pombal and Haddad, 1992 | 0 | 0 | 0 | 88 | 0 | 0 | 0 | 7 | 0 | 0 | 0 | 0 | 0 | 0 | 6 | 0 | 0 | 0 | 0 | 0 | 0 | 19 | 0 | 0 | 56 | 0 | 0 | 0 |
| RANIDAE |  |  |  |  |  |  |  |  |  |  |  |  |  |  |  |  |  |  |  |  |  |  |  |  |  |  |  |  |
| *Lithobates catesbeianus* (Shaw 1802) | 0 | 0 | 0 | 0 | 0 | 0 | 0 | 25 | 0 | 10 | 0 | 0 | 0 | 0 | 0 | 0 | 0 | 0 | 2 | 0 | 0 | 8 | 0 | 0 | 0 | 0 | 0 | 0 |
| Total abundance | 11 | 76 | 23 | 100 | 170 | 400 | 126 | 98 | 56 | 89 | 19 | 162 | 105 | 142 | 63 | 104 | 21 | 163 | 147 | 66 | 114 | 99 | 36 | 78 | 112 | 124 | 90 | 67 |
| Total richness | 2 | 2 | 3 | 3 | 3 | 1 | 3 | 8 | 2 | 5 | 4 | 5 | 6 | 2 | 6 | 2 | 2 | 2 | 6 | 2 | 3 | 7 | 2 | 2 | 4 | 2 | 1 | 1 |
| Total |  |  |  |  |  |  |  |  |  |  |  |  |  |  |  |  |  |  |  |  |  |  |  |  |  |  |  | 2,861 |

Page 43 of 60

1

2

3

4

5

6

7

8

9

10

11

12

13

14

15

16

17

18

19

20

21

22

23

24

25

26

27

28

29

30

31

32

33

34

35

36

37

38

39

40

41

42

43

44

45

46

47

48

49

50

51

52

53

54

55

56

57

58

59

60

**TABLE S2** Summary of Monte-Carlo tests of RLQ models of ponds.

|  |  | Functional | | Predictors |  |  |  |
| --- | --- | --- | --- | --- | --- | --- | --- |
| Model |  | traits |  |  |  | Std. observed | P-value |
|  |  |  |  | selected |  |  |  |
|  |  | selected | |  |  |  |  |
| Local |  |  |  |  |  |  |  |
| environmental |  | All traits | | All predictors | | 3.021 | 0.02 |
| descriptors | – |  |  |  |  |  |  |
| global model |  |  |  |  |  |  |  |
|  |  | BH; BL; BW; | | Depth; water pH; | |  |  |
|  |  | TMW; | DFH; |  |  |  |  |
| Local |  |  |  | water |  |  |  |
|  |  | VFH; | RNT; |  |  |  |  |
| environmental |  |  |  | temperature; | |  |  |
|  |  | ODP; | MS; |  |  | 3.687 | 0.001 |
| descriptors – fitted | |  |  | water |  |  |  |
|  |  | ES; SL; SW; | |  |  |  |  |
| model |  |  |  | conductivity; pond | |  |  |
|  |  | ND; ED; Fl; | |  |  |  |  |
|  |  |  |  | vegetation |  |  |  |
|  |  | EP; SP; NP | |  |  |  |  |
| Landscape |  |  |  |  |  |  |  |
| descriptors | – | All traits | | All predictors | | 1.523 | 0.06 |
| global model |  |  |  |  |  |  |  |
|  |  | BH; BL; BW; | |  |  |  |  |
|  |  | TMW; | DFH; | Forest; |  |  |  |
| Landscape |  | VFH; | RNT; | agriculture; |  |  |  |
| descriptors – fitted | | ODP; | MS; | pasture; | urban; | 1.051 | 0.15 |
| model |  | ES; SL; SW; | | edge dist.; | types |  |  |
|  |  | ND; ED; Fl; | | of land use |  |  |  |
|  |  | EP; SP; NP | |  |  |  |  |

1

2

3

4

5

6

7

8

9

10

11

12

13

14

15

16

17

18

19

20

21

22

23

24

25

26

27

28

29

30

31

32

33

34

35

36

37

38

39

40

41

42

43

44

45

46

47

48

49

50

51

52

53

54

55

56

57

58

59

60

Page 44 of 60

**TABLE S3** Pond variables and RLQ axes. Q1 = Axis 1; Q2= Axis 2; Local environment descriptor = Local physicochemical and morphological characteristics; Std. obs. values = Standard observed values; Adj. p-value = Adjusted p-value.

| Local environment | Std. obs. values | Adj. p-value |
| --- | --- | --- |
| RLQ axes |  |  |
| descriptor |  |  |
| Water depth | -2.570 | 0.006 |
| Water pH | -2.346 | 0.008 |
| Water temperature | -1.333 | 0.20 |
| Water conductivity | 1.287 | 0.20 |
| Pond grass | -0.462 | 0.32 |
| Q1 |  |  |
| Pond | 2.384 | 0.94 |
| grass+monocultures |  |  |
| Pond grass+trees | 0.401 | 0.62 |
| Pond shrubs | -0.535 | 0.38 |
| Pond trees | -0.916 | 0.17 |
| Water depth | -0.270 | 0.80 |
| Water pH | 0.576 | 0.58 |
| Water temperature | 1.336 | 0.58 |
| Water conductivity | -1.650 | 0.05 |
| Pond grass | 0.191 | 0.60 |
| Q2 |  |  |
| Pond | -0.408 | 0.5 |
| grass+monocultures |  |  |
| Pond grass+trees | 0.661 | 0.75 |
| Pond shrubs | -0.719 | 0.29 |
| Pond trees | -1.397 | 0.05 |

Page 45 of 60

1

2

3

4

5

6

7

8

9

10

11

12

13

14

15

16

17

18

19

20

21

22

23

24

25

26

27

28

29

30

31

32

33

34

35

36

37

38

39

40

41

42

43

44

45

46

47

48

49

50

51

52

53

54

55

56

57

58

59

60

**TABLE S4** Attributes (pond variables) and RLQ axes; R1 = Axis 1; R2= Axis 2; Functional traits = Measured characteristics of tadpoles; Std. obs. values = Standard observed values; Adj. p-value = Adjusted p-value.

| RLQ axes | Functional traits | Std. obs. values | Adj. p-value |
| --- | --- | --- | --- |
|  | BH | -0.382 | 0.72 |
|  | BL | 1.649 | 0.09 |
|  | BW | 2.027 | 0.05 |
|  | TMW | -1.752 | 0.05 |
|  | DFH | 2.121 | 0.02 |
|  | VFH | 0.856 | 0.44 |
|  | RNT | -0.309 | 0.75 |
|  | ODP anterior | -0.302 | 0.41 |
|  | ODP anteroventral | -0.031 | 0.47 |
|  | ODP ventral | 0.00 | 0.99 |
|  | MS | -1.777 | 0.05 |
|  | ES | -1.883 | 0.05 |
|  | SL | 0.561 | 0.62 |
|  | SW | 0.903 | 0.39 |
|  | ND | -0.991 | 0.33 |
| R1 | ED | -2.145 | 0.02 |
|  | Fl absent | -1.985 | 0.04 |
|  | Fl present | 0.759 | 0.77 |
|  | EP dorsal | -2.148 | 0.009 |
|  | EP dorsolateral | 0.110 | 0.56 |
|  | EP lateral | -0.003 | 0.43 |
|  | SP posterior | -0.528 | 0.35 |
|  | SP posterodorsal | -2.417 | 0.02 |
|  | SP sinistral | 0.000 | 0.99 |
|  | SP ventral | 0.547 | 0.75 |
|  | NP anteroventral | -0.496 | 0.38 |
|  | NP absent | 0.016 | 0.57 |
|  | NP dorsal | -0.601 | 0.33 |
|  | NP dorsolateral | -1.145 | 0.12 |
|  | NP lateral | -1.010 | 0.16 |
|  | NP laterodorsal | 0.000 | 0.99 |
|  | BH | 1.846 | 0.05 |
| R2 | BL | 0.110 | 0.92 |
|  | BW | -1.264 | 0.23 |
|  | TMW | 1.129 | 0.26 |

1

2

3

4

5

6

7

8

9

10

11

12

13

14

15

16

17

18

19

20

21

22

23

24

25

26

27

28

29

30

31

32

33

34

35

36

37

38

39

40

41

42

43

44

45

46

47

48

49

50

51

52

53

54

55

56

57

58

59

60

Page 46 of 60

|  | DFH | 1.403 | 0.17 |
| --- | --- | --- | --- |
|  | VFH | 0.112 | 0.91 |
|  | RNT | -0.913 | 0.40 |
|  | ODP anterior | 0.369 | 0.65 |
|  | ODP anteroventral | -0.763 | 0.24 |
|  | ODP ventral | 0.000 | 1.00 |
|  | MS | 1.458 | 0.15 |
|  | ES | 0.695 | 0.49 |
|  | SL | -0.315 | 0.79 |
|  | SW | 0.946 | 0.36 |
|  | ND | 1.220 | 0.23 |
|  | ED | 0.540 | 0.59 |
|  | Fl absent | -1.844 | 0.04 |
|  | Fl present | 2.127 | 0.98 |
|  | EP dorsal | -0.806 | 0.24 |
|  | EP dorsolateral | -0.325 | 0.38 |
|  | EP lateral | 0.734 | 0.75 |
|  | SP posterior | 0.178 | 0.60 |
|  | SP posterodorsal | -2.556 | 0.01 |
|  | SP sinistral | 0.000 | 1.00 |
|  | SP ventral | 1.709 | 0.95 |
|  | NP anteroventral | -0.017 | 0.61 |
|  | NP absent | 0.572 | 0.69 |
|  | NP dorsal | -1.163 | 0.11 |
|  | NP dorsolateral | -1.607 | 0.05 |
|  | NP lateral | -0.795 | 0.22 |
|  | NP laterodorsal | 0.000 | 1.00 |

Page 47 of 60

1

2

3

4

5

6

7

8

9

10

11

12

13

14

15

16

17

18

19

20

21

22

23

24

25

26

27

28

29

30

31

32

33

34

35

36

37

38

39

40

41

42

43

44

45

46

47

48

49

50

51

52

53

54

55

56

57

58

59

60

**TABLE S5** Pond landscape descriptors and RLQ axes; Q1 = Axis 1; Q2= Axis 2; Landscape descriptor = Landscape descriptor data; Std. obs. values = Standard observed values; Adj. p-value = Adjusted p-value.

| RLQ axes | Landscape | Std. obs. values | Adj. p-value |
| --- | --- | --- | --- |
|  | descriptor |  |  |
|  | Forest | -0.205 | 0.84 |
|  | Agriculture | 0.687 | 0.52 |
| Q1 | Pasture | -0.119 | 0.93 |
|  | Urban | -2.780 | 0.001 |
|  | Edge dist. | -2.621 | 0.003 |
|  | Canopy open | -1.073 | 0.12 |
|  | Forest | 1.988 | 0.03 |
|  | Agriculture | -2.594 | 0.01 |
| Q2 | Pasture | 0.553 | 0.60 |
|  | Urban | 0.647 | 0.48 |
|  | Edge dist. | 0.230 | 0.87 |
|  | Canopy open | -1.843 | 0.10 |

1

2

3

4

5

6

7

8

9

10

11

12

13

14

15

16

17

18

19

20

21

22

23

24

25

26

27

28

29

30

31

32

33

34

35

36

37

38

39

40

41

42

43

44

45

46

47

48

49

50

51

52

53

54

55

56

57

58

59

60

Page 48 of 60

**TABLE S6** Attributes (pond variables) and RLQ axes; R1 = Axis 1; R2= Axis 2; Functional traits = Measured characteristics of tadpoles; Std. obs. values = Standard observed values; Adj. p-value = Adjusted p-value.

| RLQ axes | Functional traits | Std. obs. values | Adj. p-value |
| --- | --- | --- | --- |
|  | BH | 1.397 | 0.16 |
|  | BL | -0.493 | 0.68 |
|  | BW | -1.565 | 0.14 |
|  | TMW | 1.069 | 0.32 |
|  | DFH | 0.159 | 0.89 |
|  | VFH | 0.484 | 0.62 |
|  | RNT | -0.643 | 0.64 |
|  | ODP anterior | -0.324 | 0.45 |
|  | ODP anteroventral | -0.827 | 0.28 |
|  | ODP ventral | <0.001 | 1.00 |
|  | MS | 1.397 | 0.16 |
|  | ES | 1.526 | 0.10 |
|  | SL | 0.390 | 0.58 |
|  | SW | 1.088 | 0.26 |
|  | ND | 1.295 | 0.23 |
| R1 | ED | 1.437 | 0.12 |
|  | Fl absent | 0.767 | 0.70 |
|  | Fl present | -0.729 | 0.34 |
|  | EP dorsal | 0.282 | 0.64 |
|  | EP dorsolateral | 1.252 | 0.89 |
|  | EP lateral | 1.408 | 0.14 |
|  | SP posterior | 0.014 | 0.63 |
|  | SP posterodorsal | 0.086 | 0.45 |
|  | SP sinistral | <0.001 | 1.00 |
|  | SP ventral | -0.653 | 0.30 |
|  | NP anteroventral | -0.469 | 0.48 |
|  | NP absent | -0.598 | 0.27 |
|  | NP dorsal | 0.386 | 0.62 |
|  | NP dorsolateral | -0.166 | 0.61 |
|  | NP lateral | -0.238 | 0.51 |
|  | NP laterodorsal | <0.001 | 1.00 |
|  | BH | 0.239 | 0.84 |
| R2 | BL | 0.174 | 0.86 |
|  | BW | 0.148 | 0.88 |
|  | TMW | 0.236 | 0.84 |

Page 49 of 60

1

2

3

4

5

6

7

8

9

10

11

12

13

14

15

16

17

18

19

20

21

22

23

24

25

26

27

28

29

30

31

32

33

34

35

36

37

38

39

40

41

42

43

44

45

46

47

48

49

50

51

52

53

54

55

56

57

58

59

60

|  | DFH | 2.319 | 0.02 |
| --- | --- | --- | --- |
|  | VFH | 0.681 | 0.52 |
|  | RNT | -0.051 | 0.95 |
|  | ODP anterior | -1.042 | 0.16 |
|  | ODP anteroventral | 1.539 | 0.93 |
|  | ODP ventral | <0.001 | 1.00 |
|  | MS | 0.285 | 0.79 |
|  | ES | -0.071 | 0.94 |
|  | SL | 0.877 | 0.39 |
|  | SW | 1.223 | 0.23 |
|  | ND | 0.874 | 0.40 |
|  | ED | -1.148 | 0.25 |
|  | Fl absent | -2.347 | 0.02 |
|  | Fl present | 1.603 | 0.94 |
|  | EP dorsal | -1.588 | 0.04 |
|  | EP dorsolateral | -0.688 | 0.23 |
|  | EP lateral | 1.902 | 0.98 |
|  | SP posterior | -1.370 | 0.11 |
|  | SP posterodorsal | -1.773 | 0.05 |
|  | SP sinistral | <0.001 | 1.00 |
|  | SP ventral | 0.591 | 0.74 |
|  | NP anteroventral | -0.662 | 0.38 |
|  | NP absent | -0.171 | 0.47 |
|  | NP dorsal | -1.593 | 0.05 |
|  | NP dorsolateral | -0.135 | 0.36 |
|  | NP lateral | 0.113 | 0.51 |
|  | NP laterodorsal | <0.001 | 1.00 |

1

2

3

4

5

6

7

8

9

10

11

12

13

14

15

16

17

18

19

20

21

22

23

24

25

26

27

28

29

30

31

32

33

34

35

36

37

38

39

40

41

42

43

44

45

46

47

48

49

50

51

52

53

54

55

56

57

58

59

60

Page 50 of 60

**TABLE S7** Summary of Monte-Carlo tests of RLQ models of streams.

|  | Functional | Predictors | Standard |  |
| --- | --- | --- | --- | --- |
| Model | traits |  |  | P-value |
|  |  | selected | observed |  |
|  | selected |  |  |  |
| Local |  |  |  |  |
| environmental | All traits | All predictors | 3.008 | 0.003 |
| descriptors – |  |  |  |  |
| global model |  |  |  |  |
|  |  | Area; water |  |  |
|  |  | transp.; water pH; |  |  |
|  | BH; BL; BW; | water |  |  |
| Local | TMW; DFH; | temperature; |  |  |
|  | VFH; RNT; | water |  |  |
| environmental |  |  |  |  |
|  | ODP; MS; | conductivity; diss. | 3.543 | 0.001 |
| descriptors – fitted |  |  |  |  |
|  | ES; SL; SW; | ox.; DQO; NO_2_; |  |  |
| model |  |  |  |  |
|  | ND; ED; Fl; | Total alk.; alk |  |  |
|  | EP; SP; NP | HCO_3_; stream |  |  |
|  |  | vegetation; |  |  |
|  |  | stream substrate |  |  |
| Landscape |  |  |  |  |
| descriptors – | All traits | All predictors | 1.208 | 0.11 |
| global model |  |  |  |  |
|  | BH; BL; BW; |  |  |  |
|  | TMW; DFH; |  |  |  |
| Landscape | VFH; RNT; | Canopy open; |  |  |
| descriptors – fitted | ODP; MS; |  | 0.341 | 0.35 |
|  |  | types of land use |  |  |
| model | ES; SL; SW; |  |  |  |

ND; ED; Fl;

EP; SP; NP

Page 51 of 60

1

2

3

4

5

6

7

8

9

10

11

12

13

14

15

16

17

18

19

20

21

22

23

24

25

26

27

28

29

30

31

32

33

34

35

36

37

38

39

40

41

42

43

44

45

46

47

48

49

50

51

52

53

54

55

56

57

58

59

60

**TABLE S8** Stream local descriptors and RLQ axes. Q1 = Axis 1; Q2= Axis 2; Local environment descriptor = Local physicochemical and morphological characteristics; Std. obs. values = Standard observed values; Adj. p-value = Adjusted p-value.

|  | Local environment | Std. obs. values | Adj. p-value |
| --- | --- | --- | --- |
| RLQ axes | |  |  |
|  | descriptor |  |  |
|  | Stream area | -0.916 | 0.38 |
|  | Water transparency | -0.904 | 0.39 |
|  | Water pH | -2.453 | 0.008 |
|  | Water temperature | -2.397 | 0.01 |
|  | Dissolved oxygen | -1.013 | 0.33 |
|  | Water conductivity | 2.162 | 0.01 |
|  | COD | 0.191 | 0.87 |
|  | NO_2_ | 1.358 | 0.18 |
|  | Total alk | 2.122 | 0.03 |
|  | Alk HCO_3_ | 2.122 | 0.03 |
|  | Stream grass | 0.188 | 0.81 |
| Q1 | | 1.685 | 0.88 |
|  | Stream grass+trees |  |  |
|  | Stream shrubs | -0.818 | 0.29 |
|  | Stream trees | -0.973 | 0.14 |
|  | Stream aquatic veg. | 0.933 | 0.78 |
|  | Stream grass+rocks | -0.811 | 0.16 |
|  | Stream | -0.747 | 0.31 |
|  | leaves+roots+rocks |  |  |
|  | Stream mud | 1.173 | 0.80 |
|  | Stream mud+rocks | -0.854 | 0.04 |
|  | Stream rocks | 0.000 | 1.00 |
|  | Stream roots+roocks | -0.834 | 0.20 |
|  | Stream area | -1.542 | 0.12 |
|  | Water transparency | -0.781 | 0.46 |
|  | Water pH | 1.301 | 0.20 |
|  | Water temperature | 1.314 | 0.20 |
|  | Dissolved oxygen | 0.496 | 0.65 |
|  | Water conductivity | 1.471 | 0.15 |
| Q2 | | -0.105 | 0.91 |
|  | COD |  |  |
|  | NO_2_ | 1.110 | 0.28 |
|  | Total alk | 0.612 | 0.58 |
|  | Alk HCO_3_ | 0.612 | 0.58 |
|  | Stream grass | -0.864 | 0.30 |
|  | Stream grass+trees | 1.036 | 0.90 |

1

2

3

4

5

6

7

8

9

10

11

12

13

14

15

16

17

18

19

20

21

22

23

24

25

26

27

28

29

30

31

32

33

34

35

36

37

38

39

40

41

42

43

44

45

46

47

48

49

50

51

52

53

54

55

56

57

58

59

60

Page 52 of 60

|  | Stream shrubs | 1.403 | 0.88 |
| --- | --- | --- | --- |
|  | Stream trees | -2.112 | 0.03 |
|  | Stream aquatic veg. | 0.989 | 0.85 |
|  | Stream grass+rocks | -1.087 | 0.04 |
|  | Stream | -1.078 | 0.19 |
|  | leaves+roots+rocks |  |  |
|  | Stream mud | -0.783 | 0.32 |
|  | Stream mud+rocks | 0.354 | 0.69 |
|  | Stream rocks | 0.000 | 1.00 |
|  | Stream roots+roocks | -0.980 | 0.14 |

Page 53 of 60

1

2

3

4

5

6

7

8

9

10

11

12

13

14

15

16

17

18

19

20

21

22

23

24

25

26

27

28

29

30

31

32

33

34

35

36

37

38

39

40

41

42

43

44

45

46

47

48

49

50

51

52

53

54

55

56

57

58

59

60

**TABLE S9** Attributes (stream descriptors) and RLQ axes; R1 = Axes 1; R2= Axes 2; Std. obs.

values = Standard observed values; Adj. p-value = Adjusted p-value.

| RLQ axes | Environment | Std. obs. values | Adj. p-value |
| --- | --- | --- | --- |
|  | descriptor |  |  |
|  | BH | 0.591 | 0.56 |
|  | BL | -1.873 | 0.05 |
|  | BW | -2.127 | 0.02 |
|  | TMW | 2.136 | 0.02 |
|  | DFH | -0.645 | 0.55 |
|  | VFH | 1.015 | 0.32 |
|  | RNT | 0.005 | 0.99 |
|  | ODP anterior | -0.768 | 0.03 |
|  | ODP anteroventral | 0.049 | 0.36 |
|  | MS | 1.537 | 0.11 |
|  | ES | 1.868 | 0.05 |
| R1 | SL | -1.104 | 0.28 |
|  | SW | -0.226 | 0.81 |
|  | ND | 1.754 | 0.07 |
|  | ED | 1.751 | 0.07 |
|  | Fl absent | -1.292 | 0.14 |
|  | Fl present | 0.766 | 0.74 |
|  | EP dorsal | -1.417 | 0.08 |
|  | EP dorsolateral | 1.100 | 0.87 |
|  | EP lateral | 0.789 | 0.75 |
|  | SP posterior | 0.222 | 0.67 |
|  | SP posterodorsal | -0.551 | 0.24 |
|  | SP ventral | 0.738 | 0.79 |

|  |  |  | |  | Page 54 of 60 |
| --- | --- | --- | --- | --- | --- |
| 1 |  |  |  |  |  |
| 2 |  |  |  |  |  |
| 3 |  | NP anteroventral | 0.738 | 0.79 |  |
| 4 |  |  |  |  |  |
| 5 |  | NP dorsal | -1.849 | 0.03 |  |
| 6 |  |  |  |  |  |
| 7 |  | NP dorsolateral | -0.492 | 0.35 |  |
| 8 |  |  |  |  |  |
| 9 |  |  |  |  |  |
| 10 |  | NP lateral | 0.060 | 0.60 |  |
| 11 |  |  |  |  |  |
| 12 |  | BH | -0.521 | 0.64 |  |
| 13 |  |  |  |  |  |
| 14 |  | BL | -0.591 | 0.57 |  |
| 15 |  |  |  |  |  |
| 16 |  | BW | 0.236 | 0.82 |  |
| 17 |  |  |  |  |  |
| 18 |  | TMW | -1.062 | 0.30 |  |
| 19 |  |  |  |  |  |
| 20 |  | DFH | 1.449 | 0.14 |  |
| 21 |  |  |  |  |  |
| 22 |  | VFH | 1.572 | 0.12 |  |
| 23 |  |  |  |  |  |
| 24 |  | RNT | -1.422 | 0.16 |  |
| 25 |  |  |  |  |  |
| 26 |  | ODP anterior | -0.785 | 0.23 |  |
| 27 |  |  |  |  |  |
| 28 |  | ODP anteroventral | -1.370 | 0.11 |  |
| 29 |  |  |  |  |  |
| 30 |  |  |  |  |  |
| 31 |  | MS | -0.911 | 0.38 |  |
| 32 |  |  |  |  |  |
| 33 |  | ES | -0.916 | 0.37 |  |
| 34 |  |  |  |  |  |
| 35 | R2 | SL | -1.580 | 0.12 |  |
| 36 |  |  |  |  |  |
| 37 |  | SW | 0.060 | 0.96 |  |
| 38 |  |  |  |  |  |
| 39 |  | ND | -1.342 | 0.19 |  |
| 40 |  |  |  |  |  |
| 41 |  | ED | -1.342 | 0.18 |  |
| 42 |  |  |  |  |  |
| 43 |  | Fl absent | -0.648 | 0.26 |  |
| 44 |  |  |  |  |  |
| 45 |  | Fl present | 0.982 | 0.84 |  |
| 46 |  |  |  |  |  |
| 47 |  | EP dorsal | -1.594 | 0.04 |  |
| 48 |  |  |  |  |  |
| 49 |  | EP dorsolateral | 1.440 | 0.93 |  |
| 50 |  |  |  |  |  |
| 51 |  | EP lateral | 0.811 | 0.79 |  |
| 52 |  |  |  |  |  |
| 53 |  |  |  |  |  |
| 54 |  | SP posterior | 0.182 | 0.60 |  |
| 55 |  |  |  |  |  |
| 56 |  | SP posterodorsal | -0.597 | 0.26 |  |
| 57 |  |  |  |  |  |
| 58 |  | SP ventral | 0.968 | 0.82 |  |
| 59 |  |  |  |  |  |
| 60 |  |  |  |  |  |

Page 55 of 60

1

2

3

4

5

6

7

8

9

10

11

12

13

14

15

16

17

18

19

20

21

22

23

24

25

26

27

28

29

30

31

32

33

34

35

36

37

38

39

40

41

42

43

44

45

46

47

48

49

50

51

52

53

54

55

56

57

58

59

60

| NP anteroventral | 0.986 | 0.82 |
| --- | --- | --- |
| NP dorsal | -1.925 | 0.02 |
| NP dorsolateral | 1.137 | 0.87 |
| NP lateral | -0.189 | 0.41 |

**TABLE S10** Stream landscape descriptors and RLQ axes. Q1 = Axes 1; Q2= Axes 2; Environment Descriptor = environment descriptor; Std. obs. values = Standard observed values; Adj. p-value = Adjusted p-value.

| RLQ axes | Environment | Std. obs. values | Adj. p-value |
| --- | --- | --- | --- |
|  | descriptor |  |  |
|  | Forest | -1.491 | 0.14 |
|  | Agriculture | -1.427 | 0.15 |
| Q1 | Pasture | 0.368 | 0.72 |
|  | Urban | 2.063 | 0.03 |
|  | Edge dist. | 1.796 | 0.08 |
|  | Canopy open | 1.758 | 0.08 |
|  | Forest | -0.838 | 0.42 |
|  | Agriculture | 0.808 | 0.43 |
| Q2 | Pasture | -1.646 | 0.10 |
|  | Urban | -1.238 | 0.18 |
|  | Edge dist. | -1.444 | 0.16 |
|  | Canopy open | 1.438 | 0.15 |

1

2

3

4

5

6

7

8

9

10

11

12

13

14

15

16

17

18

19

20

21

22

23

24

25

26

27

28

29

30

31

32

33

34

35

36

37

38

39

40

41

42

43

44

45

46

47

48

49

50

51

52

53

54

55

56

57

58

59

60

Page 56 of 60

**TABLE S11** Attributes (stream landscapes descriptors) and RLQ axes; R1 = Axes 1; R2= Axes 2; Environment Descriptor = environment descriptor; Std. obs. values = Standard observed values; Adj. p-value = Adjusted p-value.

| RLQ axes | Environment | Std. obs. values | Adj. p-value |
| --- | --- | --- | --- |
|  | descriptor |  |  |
|  | BH | 1.067 | 0.31 |
|  | BL | -2.236 | 0.02 |
|  | BW | -1.909 | 0.05 |
|  | TMW | 1.976 | 0.04 |
|  | DFH | -0.769 | 0.46 |
|  | VFH | 1.536 | 0.12 |
|  | RNT | -0.896 | 0.38 |
|  | ODP anterior | -0.818 | 0.09 |
|  | ODP anteroventral | -1.205 | 0.13 |
|  | MS | 1.066 | 0.31 |
|  | ES | 1.792 | 0.07 |
|  | SL | -1.989 | 0.03 |
|  | SW | -0.646 | 0.55 |
| R1 | ND | 1.815 | 0.05 |
|  | ED | 1.689 | 0.08 |
|  | Fl absent | -1.100 | 0.14 |
|  | Fl present | -0.514 | 0.27 |
|  | EP dorsal | -1.553 | 0.08 |
|  | EP dorsolateral | 0.784 | 0.78 |
|  | EP lateral | -0.494 | 0.27 |
|  | SP posterior | 1.917 | 0.95 |
|  | SP posterodorsal | -1.756 | 0.04 |
|  | SP ventral | -0.358 | 0.42 |
|  | NP anteroventral | -0.358 | 0.42 |
|  | NP dorsal | -1.928 | 0.03 |
|  | NP dorsolateral | -0.648 | 0.28 |
|  | NP lateral | -0.215 | 0.40 |
|  | BH | 0.057 | 0.96 |
|  | BL | 0.031 | 0.98 |
|  | BW | -0.199 | 0.85 |
| R2 | TMW | 1.729 | 0.07 |
|  | DFH | -0.823 | 0.41 |
|  | VFH | -0.861 | 0.40 |
|  | RNT | 2.187 | 0.02 |

Page 57 of 60

1

2

3

4

5

6

7

8

9

10

11

12

13

14

15

16

17

18

19

20

21

22

23

24

25

26

27

28

29

30

31

32

33

34

35

36

37

38

39

40

41

42

43

44

45

46

47

48

49

50

51

52

53

54

55

56

57

58

59

60

|  | ODP anterior | -0.737 | 0.12 |
| --- | --- | --- | --- |
|  | ODP anteroventral | -0.791 | 0.18 |
|  | MS | 1.834 | 0.05 |
|  | ES | 1.390 | 0.18 |
|  | SL | 0.922 | 0.38 |
|  | SW | -0.516 | 0.64 |
|  | ND | 1.612 | 0.11 |
|  | ED | 1.619 | 0.10 |
|  | Fl absent | -1.435 | 0.09 |
|  | Fl present | 1.626 | 0.92 |
|  | EP dorsal | -1.608 | 0.02 |
|  | EP dorsolateral | 1.488 | 0.93 |
|  | EP lateral | 1.627 | 0.93 |
|  | SP posterior | -0.542 | 0.39 |
|  | SP posterodorsal | -0.725 | 0.24 |
|  | SP ventral | 2.099 | 0.95 |
|  | NP anteroventral | 2.099 | 0.95 |
|  | NP dorsal | -1.689 | 0.03 |
|  | NP dorsolateral | 0.137 | 0.62 |
|  | NP lateral | -1.274 | 0.05 |

1

2

3

4

5

6

7

8

9

10

11

12

13

14

15

16

17

18

19

20

21

22

23

24

25

26

27

28

29

30

31

32

33

34

35

36

37

38

39

40

41

42

43

44

45

46

47

48

49

50

51

52

53

54

55

56

57

58

59

60

Page 58 of 60

Figure S1. The photographic images show details of the landscape of each of the waterbodies sampled in the tadpole collection, in the areas of Atlantic Forest, in southern Brazil.

Page 59 of 60

1

2

3

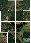

4

5

6

7

8

9

10

11

12

13

14

15

16

17

18

19

20

21

22

23

24

25

26

27

28

29

30

31

32

33

34

35

36

37

38

39

40

41

42

43

44

45

46

47

48

49

50

51

52

53

54

55

56

57

58

59

60

Page 60 of 60

1

2

3

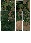

4

5

6

7

8

9

10

11

12

13

14

15

16

17

18

19

20

21

22

23

24

25

26

27

28

29

30

31

32

33

34

35

36

37

38

39

40

41

42

43

44

45

46

47

48

49

50

51

52

53

54

55

56

57

58

59

60
